# Somatic epigenetic drift during shoot branching: a cell lineage-based model

**DOI:** 10.1101/2024.01.24.577071

**Authors:** Yifan Chen, Agata Burian, Frank Johannes

## Abstract

Plant architecture is shaped by the continuous production of new organs, most of which emerge post-embryonically. This process includes the formation of new lateral branches along existing shoots. Shoot branching is fundamental to plant development, plant-environment interactions, and vegetative propagation. Current empirical evidence supports a “detached meristem” model as the cellular basis of lateral shoot initiation. In this model, a small number of undifferentiated cells are “sampled” from the periphery of the shoot apical meristem (SAM) to act as precursors for axillary buds, which eventually develop into new shoots. Repeated branching thus creates a series of cellular bottlenecks (i.e. somatic drift) that affect how *de novo* genetic and epigenetic mutations propagate through the plant body during development. Somatic drift could be particularly relevant for epigenetic changes in the form of stochastic DNA methylation gains and losses (i.e. spontaneous epimutations), as they have been shown to arise rapidly with each cell division.

Here, we formalize a special case of the “detached meristem” model, where precursor cells are randomly sampled from the SAM periphery in a way that maximizes cell lineage independence. By following a population of SAM cells through repeated branching processes, we show that somatic drift gives rise to a complex mixture of cellular phylogenies, which shape the evolution of cell-to-cell DNA methylation heterogeneity within the SAM over time. This process is dependent on the number of branch points, the strength of somatic drift as well as the epimutation rate. For many realistic cell biological settings, our model predicts that cell-to-cell DNA methylation heterogeneity in the SAM converges to non-zero states during development, suggesting that epigenetic variation is an inherent property of the SAM cell population.

Our insights have direct implications for empirical studies of somatic (epi)genomic diversity in long-lived perennial and clonal species using bulk or single-cell sequencing approaches.

## 1 Introduction

In contrast to most animals, plants have the ability to undergo continuous organogenesis throughout their entire lifetime. A notable example is shoot branching, whereby new lateral shoots are formed repeatedly along an existing shoot, often giving rise to elaborate post-embryonic body plans (**Fig. 1A**). Thus, branching is essential for development as it determines the overall architecture of the plant body, enabling the exploration of space, light capture, and an increase in the number of leaves, flowers or fruits. In addition, it plays a crucial role in vegetative propagation, such as in the formation of runners, rhizomes, or tillers. The plasticity of shoot branching, demonstrated through a vast diversity of branching patterns and strategies, allows plants to respond and adopt to changing environment (Domagalska and Leyser, 2011). The cellular and molecular mechanisms underlying shoot branching have been well characterized.

**Figure 1:**
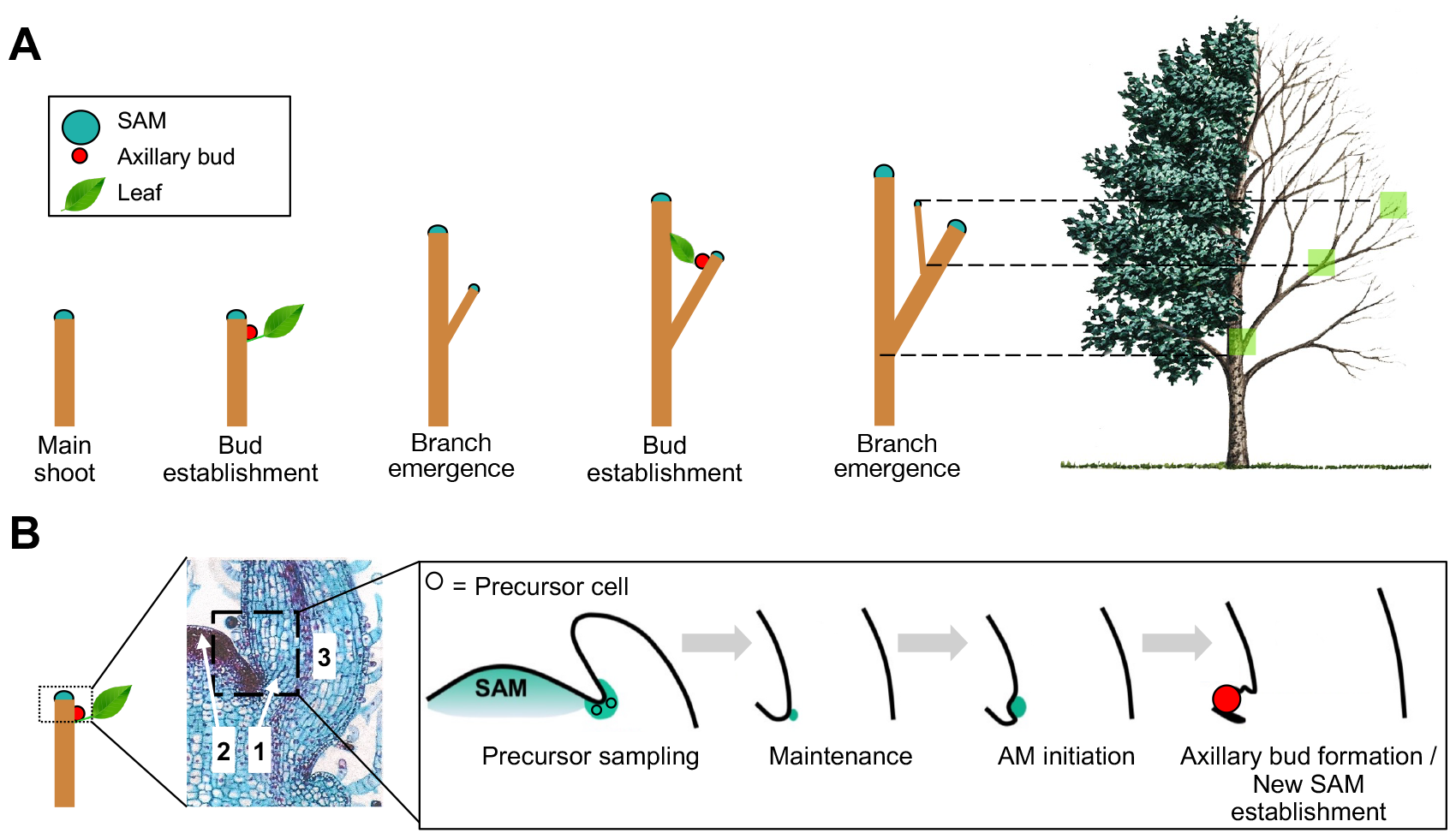
Shoot branching and the detached-meristem model: **A**. New lateral branches emerge from axillary buds localized at the leaf axils of the main shoot. This process repeats on newly formed branches, eventually giving rise to complex plant architectures, like those observed in trees. **B**. According to the detached-meristem model, the axillary bud develops from the axillary meristem (AM), which originates from a few precursor cells sequestered from the shoot apical meristem (SAM). These precursors are maintained in an undifferentiated state in the leaf axil until the activation of AM initiation, followed by the formation of the axillary bud with a newly established SAM. (B-1: Location of the AM in the leaf axil, B-2: Shoot apical meristem, B-3: Emerging leaf). Figure 1B is modified from Yang et al., 2023.

### 1.1 The cellular basis of shoot branching

In higher plants, lateral branches are initiated from axillary buds, which are formed at the axils of developing leaves along the stem (**Fig. 1A, B**). The buds, themselves, derive from a cluster of cells, known as axillary meristems (AM) (**Fig. 1B**). The cellular origin of AMs has been subject of debate (McSteen and Leyser, 2005). However, recent studies using live imaging support a “detached-meristem” model, where AMs derive from a few precursor cells sequestered from the periphery of the shoot apical meristem (SAM) (Burian et al., 2016; Shi et al., 2016). These precursors maintain their meristematic characteristics in the leaf axil, eventually increase in number and develop into active AMs generating new organs. As such they become the SAM of the newly emerging shoot, and share all the functional and morphological properties with the original SAM (e.g. organization, capability of self-maintenance and organ formation (Nicolas and Laufs, 2022)). Thus, shoot branching represents a developmental transition from a SAM at the original shoot to the establishment of a new SAM on the emerging lateral branch.

The SAM itself is characterized by a tunica-corpus structure, which refers to the arrangement of cells in the outer layers (tunica) and the inner region (corpus). In most plant species, the tunica consists of one or two cell layers (L1, L2) where cells divide anticlinally (perpendicular to the surface), while in the underlying corpus (L3), cells divide in various directions (Lyndon, 1998). The SAM consists of a few hundreds of cells, and its function is not only related to the generation of new organs, but also to self-maintenance, enabling restoration of the cell pool “used” for organ formation (Kuhlemeier, 2017). Crucial for its function, the SAM contains stem cells with the ability to renew and to generate cells that enter the differentiation pathway (Heidstra and Sabatini, 2014). Live imaging and cell lineage tracing have revealed 3-4 stem cells at the very tip of the SAM surface in Arabidopsis and tomato (Burian et al., 2016). These apical stem cells can be distinguished from the surrounding cells by their extraordinary slow division rates (Kwiatkowska, 2008; Lyndon, 1998). Despite their low number and mitotic activity, apical stem cells are the ultimate source of all cell lineages that emerge within the SAM, including those that give rise to AMs as well as the germline (Burian, 2021). Thus, the population of apical stem cells (here denoted as ASC) presents an important unit for cell lineage-based analyses.

### 1.2 Somatic genetic and epigenetic drift

The sampling of AM precursors presents a major cellular bottlneck during shoot branching. It creates somatic genetic drift (Reusch et al., 2021; Yu et al., 2020), which affects how *de novo* genetic mutations that occasionally arise in the SAM are propagated through the plant body. Indeed, due to somatic drift, these mutations can rapidly increase in frequency in newly formed lateral branches. This phenomenon explains the existence of genetic mosaics observed in various plant species, where *de novo* mutations have become fixed in some shoot sectors or in whole organs but not in others (Frank and Chitwood, 2016). It has been argued that such fixed somatic mutations could become a source of within-organism or within-clone selection in modular organisms, as each module can potentially function in a (semi)automonous manner (Herrera et al., 2021; Reusch et al., 2021; Whitham and Slobodchikoff, 1981). Although observations of this selection remain scant in natural settings, fixed somatic mutations have been artificially selected for centuries in the form of phenotypically distinct shoots (i.e. bud sports) in numerous horticultural varieties, e.g. in fruit trees (Foster and Aranzana, 2018).

Nonetheless, SAM-derived genetic mutations are expected to be rare. They occur at a rate of about 10^−10^ per site per year (Hofmeister et al., 2020; Plomion et al., 2018; Satake et al., 2023; Schmid-Siegert et al., 2017). This low rate per unit time has been partly attributed to the slow mitotic division rate observed in the ASC population (Kwiatkowska, 2008; Laufs et al., 1998; Reddy et al., 2004). Similar low division rates have been found during sequestration of AM precursors (Burian et al., 2016). These observations are in line with studies in the annual plant A. thaliana, where the gamete-to-gamete cell depth has been estimated to be around only 34 mitoses (Watson et al., 2016), even though the gametes are established rather late in development. Interestingly, a similar cell depth probably exists in long-lived trees, as the *per generation* somatic mutation rate is similar to that in A. thaliana, although the *per unit of time* (e.g. per year) rate is much lower (Shahryary et al., 2020). One can speculate that the slow cell division rate of the ACS is an evolved property to alleviate the build-up of mutational load when germlines are segregated late in development (Berger and Twell, 2011; Burian, 2021; Paszkowski and Grossniklaus, 2011).

In contrast to somatic genetic mutations, the rate of somatic epigenetic changes in the form of spontaneous gains and losses of DNA methylation at CG dinucleotides is orders of magnitude higher per cell division (Johannes and Schmitz, 2019). Current molecular evidence suggests that these so-called spontaneous “epimutations” are the result of enzymatic failures of DNA methyltransferases to maintain CG methylation patterns during DNA replication (Briffa et al., 2023). Once acquired, CG epimutations can be stably inherited across mitosis. Similar to somatic genetic mutations, their propagation through the plant body is therefore subject to the same cellular sampling constraints outlined above. However, because of their high rate, epimutations accumulate more rapidly and thus constitute an abundant source of somatic cellular heterogeneity.

### 1.3 Conceptual framework

The functional significance of somatic epigenetic heterogeneity in plants remains unclear. However, a formal analysis of the post-embryonic emergence of this heterogeneity is crucial for understanding how cell-to-cell epigenetic variation in the SAM is shaped by somatic drift. This is particularly important, given that the SAM, especially the ASC population, is the ultimate source for almost all above-ground plant organs. A number of mathematical models have been introduced to study the influence of different aspects on somatic mutation accumulation, such as the cellular organization and dynamics of the SAM (Burian et al., 2016; Iwasa et al., 2023; Klekowski and Kazarinova-Fukshansky, 1984a; Klekowski et al., 1985; Pineda-krch and Lehtilä, 2002), tree development (Klekowski et al., 1989; Tomimoto and Satake, 2023; Yu et al., 2023), intra-organismal selection on mutant stem cells and modules (Antolin and Strobeck, 1985; Folse and Roughgarden, 2012; Klekowski and Kazarinova-Fukshansky, 1984b; Otto and Orive, 1995; Pineda-Krch and Fagerström, 1999), and branching architecture (Tomimoto et al., 2023). Only recently has an analysis of cell lineages been considered in mathematical models studying the SAM (Iwasa et al., 2023). Iwasa et al., 2023 employed coalescent approaches to model cell lineage phylogenies arising from different stem cell dynamics during development of a single shoot. However, in their model, neither shoot branching nor the epigenetic processes occurring in cell lineages have been considered. Here we present a cell lineage-based model that takes these factors into account.

Our model builds on the conceptual framework introduced by Tomimoto and Satake, 2023. In this framework the cellular processes between two sequential branch points are partitioned into two phases: (1) ASC self-renewal at the SAM center (corresponding to “elongation” in the model of Tomimoto and Satake, 2023), which consists of 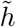 asymmetric stem cell divisions (**Fig. 2A**), and (2) branching, which consists of *h* symmetric cell divisions occurring in the peripheral zone of the SAM, followed by precursor sampling at the SAM periphery (**Fig. 2B**). We note that the symmetric cell divisions in the second phase generate spatially defined clonal sectors at the SAM surface that have their origin in each of the apical stem cells at the center (Burian et al., 2016) (**Fig. 2B**).

**Figure 2:**
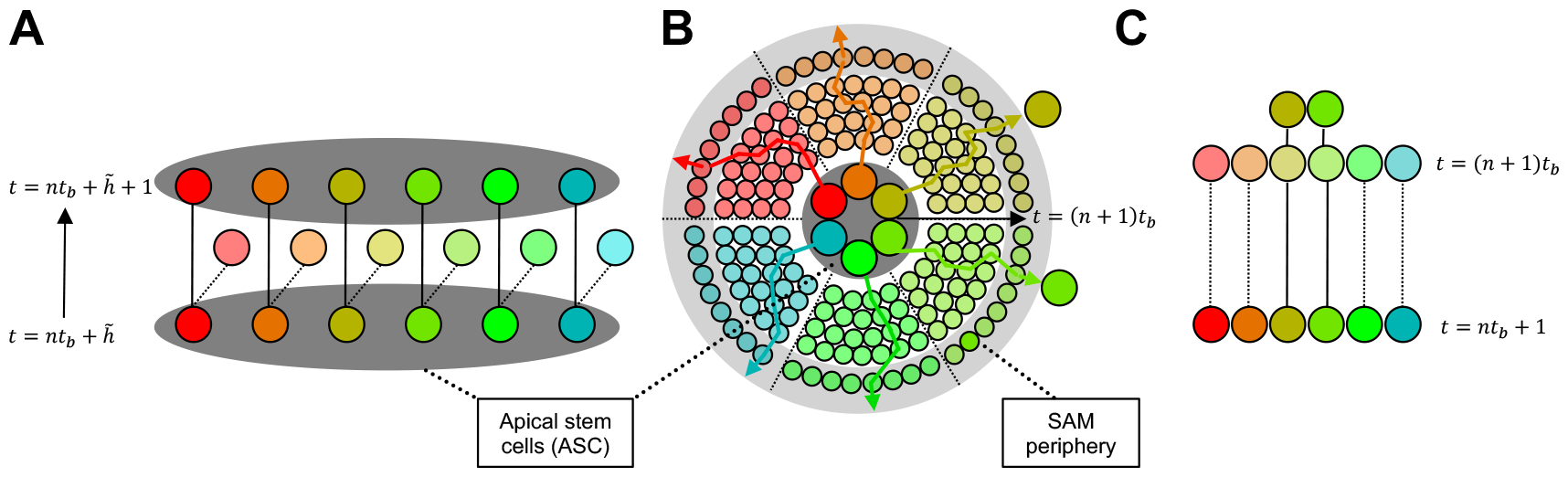
Cellular events in a cell lineage-based model of shoot branching: The cellular events between two sequential branch points at t = *nt*_*b*_ and *t* = *n +* 1 *t*_*b*_, where *n* ≥ 1 and 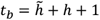, can be partitioned into a “self-renewal” and a “branching” part (Tomimoto and Satake, 2023) **A**. ASC self-renewal: A population of 6 ASCs at the SAM center (dark gray) is shown. After precursor sampling, the ASC population size is reestablished between *t* = *nt*_*b*_ and *t* = *nt*_*b*_ *+* 1, followed by 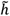 asymmetric stem cell divisions, where one descendant cell retains stem cell function and the central position, while another cell (light shaded) is displaced into the SAM periphery (light gray). From 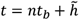 to 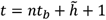, each cell of the ASC population divides one last time before the branching process begins. **B**. Branching: We trace the cell being displaced into the periphery from (A) and track only one of the descendent cells after each mitotic cell division until after another *h* mitotic divisions, the cell has “reached” the periphery. Shown is a top view of the SAM L1 layer with the ASC at the center and the SAM periphery at the edges. At time *t* = *n +* 1 *t*_*b*_, precursor sampling starts after the formation of cell lineages originating from the ASC. These cells “reached” the SAM periphery after a total of *t*_*b*_ mitotic divisions since the last branch point at *t* = *nt*_*b*_. This process generates spatially defined clonal sectors at the SAM surface (shared color sectors). Each sector has its origin in one of the ASCs. In an extreme case, which we assume in our model, sampling of precursors at the SAM periphery occurs randomly and independent across the sectors (with maximal one cell being sampled per sector). **C**. Our cell-lineage model: In the above special cases, the self-renewal and branching processes in (A) and (B) can be collapsed into a single lineage model.

We consider two special cases of self-renewal and branching. For self-renewal (phase 1), we assume that after precursor sampling at the SAM periphery, the population size of the ASC is reestablished in only one round of cell divisions, followed by 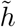 asymmetric cell divisions of the ASC (**Fig. 2A**). Given that the same population of ASCs can function for a relatively long period (Burian et al., 2016; Lyndon, 1998; Stewart and Dermen, 1970), we further assume that ASCs are not subject to turnover. For the precursor sampling during branching (phase 2), we suppose that precursor cells at the SAM periphery are selected in such a way that the clonal sectors are randomly and uniformly sampled without replacement (**Fig. 2B**). This sampling strategy neglects known spatial biases during precursor selection and thus eliminates the possibility that precursor lineages have a recent common origin in the same clonal sector of the SAM. Under these special cases, the self-renewal and branching phases can be simplified into a single cell-lineage based model of shoot branching (**Fig. 2C**). For clarity, this model measures time in units of cell divisions. Hence, a “branch point” is defined as the point in time where precursor sampling occurs, and the time segment between two sequential branch points, which we denote as *t*_*b*_, is the cumulative number of asymmetric and symmetric cell divisions arising during the self-renewal and branching phases, i.e. 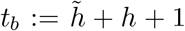. Based on empirical observations, we assume that two sequential branch points are separated by only 10-12 cell divisions (Burian et al., 2016; Romberger et al., 2004).

The importance of a cell lineage-based model for studying shoot branching is motivated in **Fig. 3**. As can be seen there, the topology and depth of cellular phylogenies that arise during sequential branching events is profoundly shaped by the size of the ASC population (*N*) as well as the strength of somatic drift (ie. the number of selected AM precursors (*M*)) (**Fig. 3A, B**). In one extreme, a population of ASCs at the SAM of newly formed lateral shoot is derived from a single precursor (*M* = 1). In this case, all these ASCs share the same “most recent common ancestor” (MRCA) cell at the last branch point, displaying maximal somatic drift and relatively low phylogenetic depth. At the other extreme, each precursor originally derives from a separate ASC at the last branch point (*M* = *N*). In this case, the MRCA theoretically traces back to the original embryonic SAM founder cell, resulting in no genetic drift and high phylogenetic depth. Any intermediate sampling scenario (1 *< M < N*) will lead to intermediate drift and non-trivial mixtures of cellular phylogenies that vary both in topology and depth. Assuming that somatic epimutations accumulate in cell lineages during successive shoot branching, different cell phylogenies will result in complex patterns of DNA methylation hetorogeneity (**Fig. 3C**). An intermediate sampling scenario is likely the most common in plants, although the precise number of precursors depends on the species and the cellular mechanisms of shoot development.

**Figure 3:**
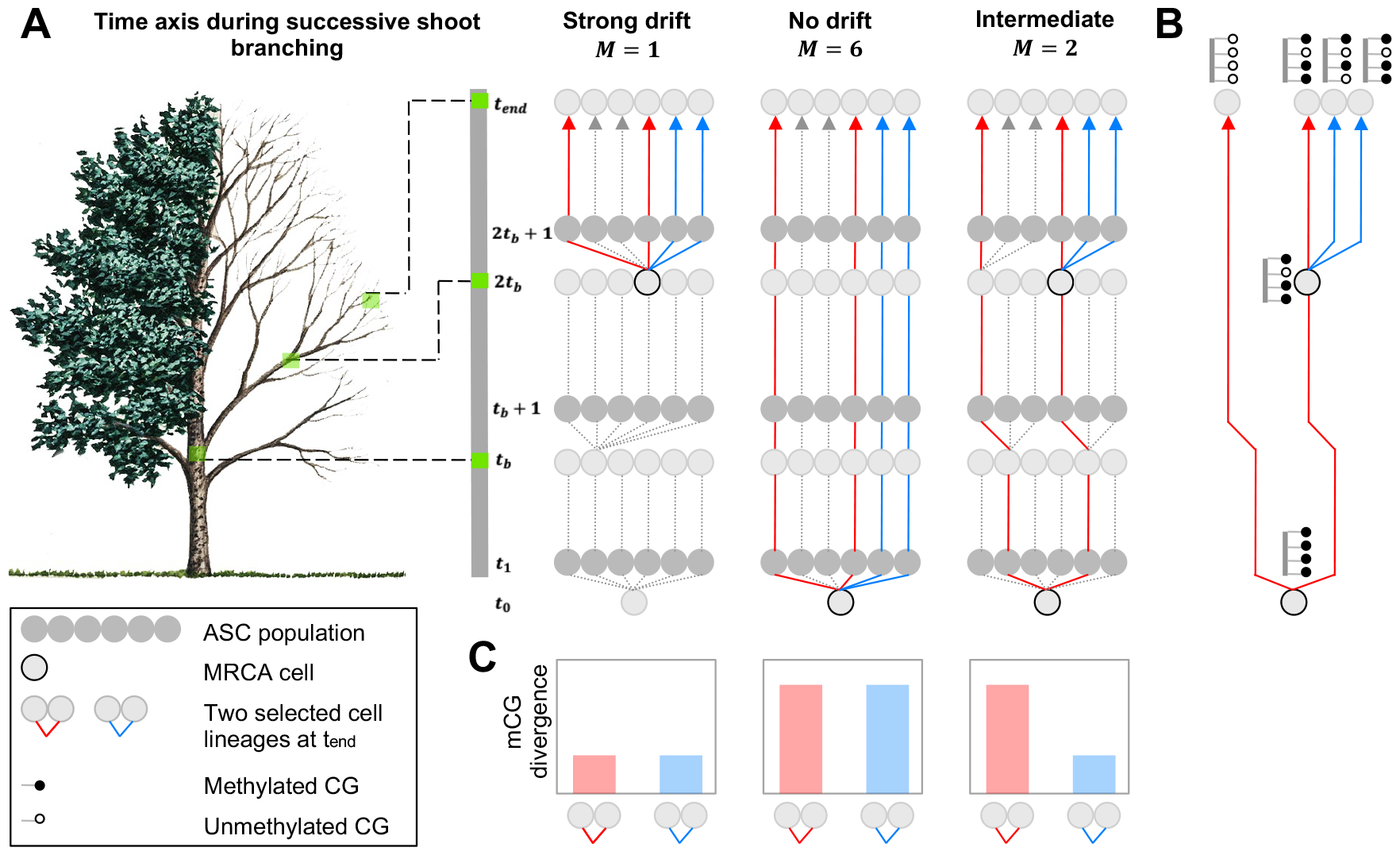
Mixtures of cell phylogenies and cell-to-cell DNA methylation heterogeneity: **A**. Schematic representations of cell lineages with different somatic drift scenarios during successive shoot branching. Branch points are indicated as *t*_*b*_and 2*t*_*b*_. The number of ASCs (*N*) is set to 6. In one scenario, drift is at its maximum with only one precursor selected at each branch point (*M* = 1). In the other scenario, drift is absent, with each ASC contributing one precursor (*M* = *N*). Although the latter scenario is biologically unlikely, it can nevertheless yield interesting theoretical insights. In both scenarios, any two selected cells from the ASC population have the same phylogenies in terms of topology and depth. However, any intermediate drift scenarios where *N* > *M* > 1 precursors are selected (here *M* = 2) can generate mixtures of cell phylogenies (blue vs. red). **B and C**. Somatic epimutations accumulate along the cell phylogenies. A mixture of cell phylogenies leads to complex cell-to-cell DNA methylation heterogeneity patterns, with cells that share a more recent common ancestor (MRCA) cell being more “similar” (low mCG divergence) than cells that share a more distal MRCA cell (high mCG divergence).

## 2 Theory

### 2.1 Sampling of precursor cells during successive branching

We focus on modeling the outermost meristematic cell layer (L1), where cellular processes are best recognized (Barbier de Reuille et al., 2015; Kwiatkowska, 2008; Lyndon, 1998). However, given the similarity of cell growth and division dynamics between layers (Jackson et al., 2019), the model output can be also applied to internal cell layers. Let the ASC of the L1 layer be defined by a population of *N* cells (*N* ∈ ℕ). We suppose that this population is initially generated from a single embryonic progenitor cell at *t* = *t*_0_. Let *t*_*b*_ denote the number of cell divisions between two sequential branch points. That is, *t*_*b*_ is the sum of the cell divisions occurring during the processes self-renewal and branching (see above). We require this number to be larger than *h* ∈ ℕ)., where *h* ≥ 1 gives the number of mitotic divisions from the last asymmetric cell division of the ASC to the SAM periphery (see **Fig. 2**). Furthermore, we assume that *t*_*b*_ is constant between two sequential branch points. Without loss of generality, we can set *t*_0_ = 0, since both the epimutation and the branching processes are constructed in such a way that they are time invariant (see below sections).

In particular, we have that for any *n* ≥ 1, between *t* = *nt*_*b*_ and *t* = *nt*_*b*_ + 1, *M* of the *N* cells are randomly selected as precursors to initiate new lateral shoots, where we assume *N/M* =: *k* ∈ ℕ).. Each of the precursor cells generates *k* daughter cells, so that the total size of the ASC remains constant over time. **Fig. 3** shows schematic representations of the two extreme sampling scenarios (*N* = 6, *M* = 1 and *N* = 6, *M* = 6) and an intermediate scenario (*N* = 6, *M* = 2), where sequential branching generates a phylogeny of cell lineages over time. The lineages of any two cells *i* and *j* can therefore be traced back to a most recent common ancestor (MRCA) cell, which is the founder of the two lineages at a past branching point.

### 2.2 Probability distribution of time to MRCA of two apical stem cells

To derive the probability distribution of the time to the MRCA of two randomly selected apical stem cells at time *t*_end_ ≥ *t*_*b*_ + 1, consider two cells *i, j*. Note that there is a total of 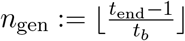 branch points up to time *t*_end_. In particular, for 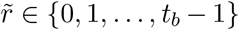, we can write *t*_end_ = *n*_gen_*t*_*b*_ + *r*, where 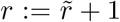. The parameter *r* describes the number of mitotic cell divisions since the last branch point, which took place at time *n*_gen_*t*_*b*_.

Further, let *t*_*i,j*_ denote the time of the MRCA of cells *i* and *j* and consider 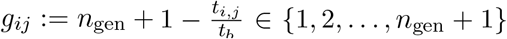. Observe that *g*_*ij*_ denotes the number of branch points to the MRCA of cells *i* and *j*, where *g*_*ij*_ = *n*_gen_ + 1 implies that the MRCA traces back to the initial progenitor cell at time *t* = 0. We have

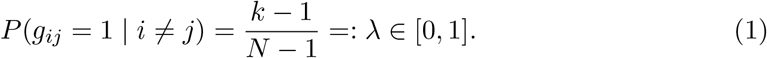

In particular, *λ* gives the probability of two cells having their MRCA at the most recent branch point.

We observe that for *M* = *N, λ* = 0 holds and for *M* = 1, we have *λ* = 1. Further, for *n* ∈ {2, 3, …, *n*_gen_}, we have

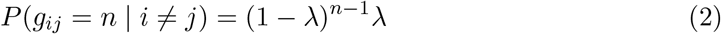

and for *n* = *n*_gen_ + 1, we have

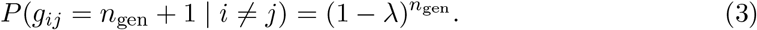

A detailed derivation of the above equations is given in the Appendix.

The above derivations reveal that the final ASC population at the last shoot is characterized by a mixture of different cellular phylogenies that differ from each other by when they share their MRCA. The mixing probabilities of these phylogenies is given by the probability distribution *P* (*g*_*ij*_ | *i* ≠*j*).

### 2.3 Epimutational processes during successive branching

As mentioned previously, in parallel to the branching processe described above, we assume that spontaneous epimutations arise at every mitotic cell division. We suppose that these epimutations are selectively neutral and that they occur independently across CG sites and homologous chromosomes. Following the work of (Shahryary et al., 2020; van der Graaf et al., 2015) we describe the epimutational process using discrete time Markov chains.

#### 2.3.1 Diploid model

Consider a locus *X*_*t*_ = (*μ, ν*) at time *t*, where *μ, ν* ∈ {*u, m*} denote the methylation state with *m* representing a methylated epiallele. Further, let *α, β* ≥ 0 denote the spontaneous gain and loss rate per site, respectively, where we assume *α* and *β* to be sufficiently small such that 0 *<* 1 − *α* − *β <* 1. Note that we do not differentiate between the states (*u, m*) and (*m, u*). For a diploid system, which is propagated clonally, 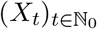 is a Markov chain with state space *S* = {(*u, u*), (*u, m*), (*m, m*)}. For the transition matrix *G*, we identify the states (*u, u*), (*u, m*), (*m, m*) with 1, 2, and 3, respectively. In particular, *G*_*i,j*_ = *P* (*X*_*t*+1_ = *j* | *X*_*t*_ = *i*). In accordance with (Shahryary et al., 2020), the transition matrix *G* is given by

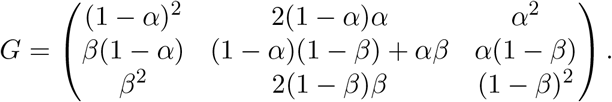

Further, let *π*(0) = (*π*(0)_1_, *π*(0)_2_, *π*(0)_3_) denote the initial distribution of *X*_0_. It holds that *π*(*t*) = (*π*(*t*)_1_, *π*(*t*)_1_, *π*)(*t*)_1_) = *π*(0) · *G*^*t*^. The stationary distribution is given by 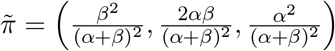.

#### 2.3.2 Haploid model

In a diploid system undergoing clonal (i.e. somatic) propagation, the two homologous chromosomes can be treated independently, and it is therefore sufficient to work with a haploid model. On a haploid chromosome, consider a locus *X*_*t*_ ∈ {*u, m*} at time *t*, which denotes the methylation state with *m* representing a methylated epiallele. Similar as before, *α, β* ≥ 0 denote the spontaneous gain and loss rate per site, respectively and we assume *α* and *β* to be sufficiently small such that 0 *<* 1 − *α* − *β <* 1. In addition, we identify the states *u, m* with 1 and 2, respectively. With *G*_*i,j*_ = *P* (*X*_*t*+1_ = *j* | *X*_*t*_ = *i*), the transition matrix *G* is given by

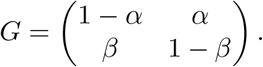

For *t >* 0, we have

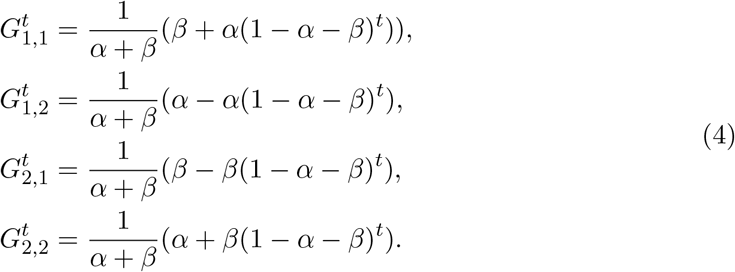

With *π*(0) = (*π*(0)_1_, *π*(0)_2_) denoting the initial distribution of *X*_0_, it holds that *π*(*t*) = (*π*(*t*)_1_, *π*(*t*)_2_) = *π*(0) · *G*^*t*^, where

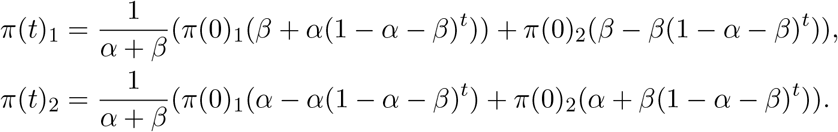

The stationary distribution is given by 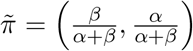.

### 2.4 Quantifying cell-to-cell DNA methylation heterogeneity

The accumulation of somatic epimutations in each cell lineage generates epigenetic heterogeneity within the ASC over time. The evolution of this heterogeneity is not only influenced by the somatic epimutation rate, but also by the strength of somatic drift arising from the finite sampling of precursor cells at each branch point. To model the evolution of cell-to-cell heterogeneity within the ASC, we derive an epigenetic distance function, which takes into account the contributions of these factors. To that end, we set aside any technical difficulties, for the moment, in obtaining haplotype-resolved DNA methylation measurements at the single-cell level.

Suppose we seek to compare the methylation states of *L* CG sites of the same haploid genome among pairs of the *N* apical stem cells. Let *t*_end_ ≥ *t*_*b*_ + 1 and indices *i* and *j* denote the same haploid genome in cells *i* and *j*, where *i* ≠ *j*. With *N* apical stem cells there are 2*N* (*N* − 1)*/*2 = *N* (*N* − 1) pairs of cells to compare. Now, consider the haploid model, where 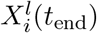 denotes the methylation state of the haploid genome in cell *i* at locus *l* at time *t*_end_. For every CG site *l* ∈ {1, …, *L*}, we consider 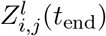, where

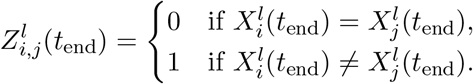

Let 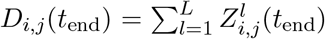, where *D*_*i,j*_(*t*_end_) ∈ {0, 1, …, *L*}. We refer to this as the cell-to-cell epigenetic distance, which gives the number of CG sites at which two cells *I* and *j* differ with respect to their methylation state. In particular, for *d* ∈ {0, 1, …, *L*} we have

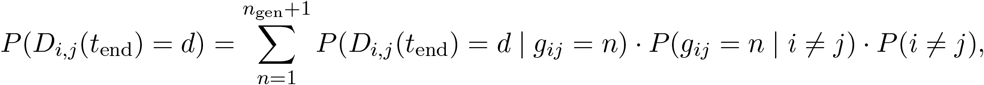

where *P* (*i* ≠ *j*) = 1. For the rest of the work, we will write *D*_*i,j*_ for simplicity.

Since we assumed the epimutational process between loci to be independent, *D*_*i,j*_ conditioned on *g*_*ij*_ = *n* follows a Binomial distribution with success probability *p*_*n*_ for *n* ∈ {1, 2, …, *n*_gen_ + 1}. Using the assumption that the initial distribution of methylation states of the loci is the stationary distribution, in the Appendix, we show that

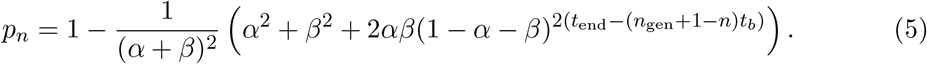

Note that we assume *α* and *β* to be sufficiently small such that 0 *<* 1 − *α* − *β <* 1. For a fixed value of 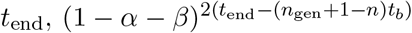 decreases for increasing values of 1 ≤ *n* ≤ *n*_gen_ + 1. On the other hand, for a fixed value of *n*, an increase in *t*_end_ leads to a decrease in 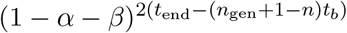. Thus, in both cases, *p*_*n*_ increases.

It follows that

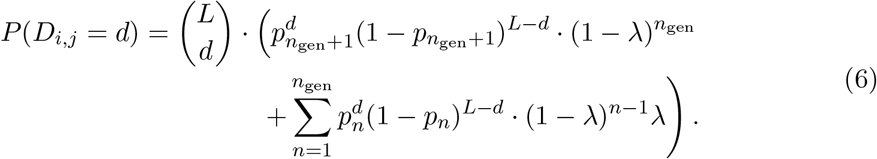

Equation (6) shows that *D*_*i,j*_ follows a multi-modal distribution, where the number of modes depends on the total number of branch points and the number of selected precursors at each branch point. For 1 *< M < N*, the number of modes is given by *n*_gen_ + 1. For *M* = 1 or *M* = *N, D*_*i,j*_ follows a single Binomial distribution. In particular, the modes are given by ⌊(*L* + 1)*p*_*n*_⌋, where the probability mass function evaluated at the modes depends on *M*.

Moreover, using the Law of total expectation and the Law of total variance, the expectation and variance of the cell-to-cell epigenetic distance are given by

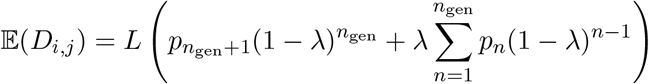

and

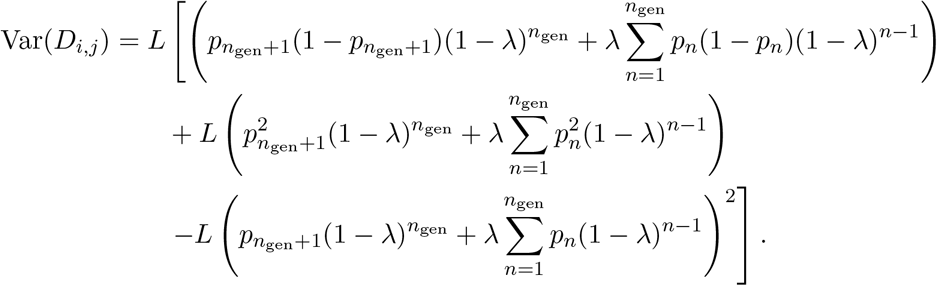

## 3 Results

### 3.1 Somatic drift shapes cell-to-cell DNA methylation heterogeneity

The sampling of precursor cells at the SAM periphery can be a major source of somatic drift. Recall that 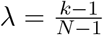 depends on the number of selected precursor cells *M* at each branching point, where 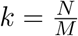. For *N* ≥ 2, the function 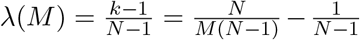 is monotone decreasing on the interval [1, *N* ] with its maximum obtained for *M* = 1. Thus, by letting all other parameter values fixed, an increase in *M* leads to a decrease in *λ* until for *M* = *N, λ* = 0 holds. We further observe that for 1 *< M < N, λ* ∈ (0, 1) holds true and thus, for a fixed value of *λ* ∈ (0, 1), *λ*(1 − *λ*)^*n*−1^ decreases with increasing *n*.

Somatic drift is strongest for *M* = 1. In this case, only one precursor is sampled from the periphery of the SAM to form the ASC of the newly emerging shoot, and (6) reduces to

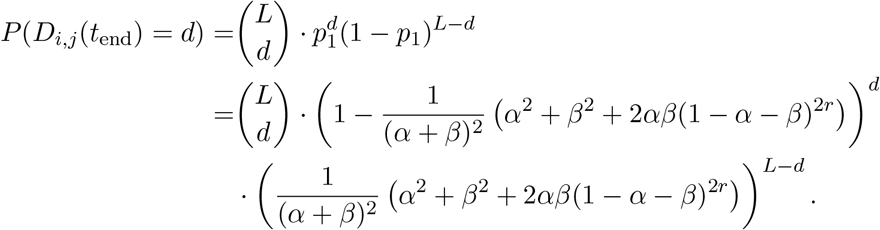

From this equation it becomes clear that the number of branch points (*n*_gen_) does not play a role in this case, since the entire process “restarts” after each branching point.

The other extreme is when drift is absent (*M* = *N*). In this case, each ASC of the branching shoot contributes to one ASC in the newly formed shoot. Thus, equation (6) reduces to

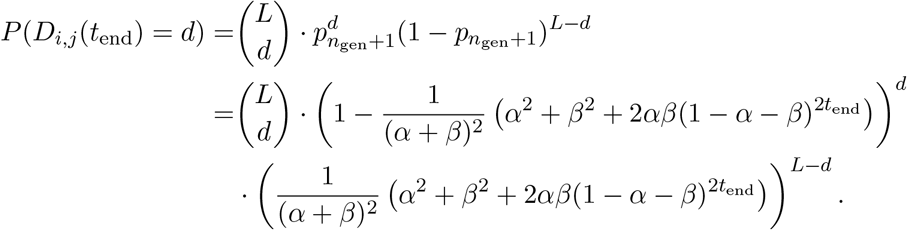

In this case, too, the branching process has no effect on the cell-to-cell epigenetic distance.

An important and perhaps less intuitive scenario arises when somatic drift is moderate; that is, when 1 *< M < N*. In this case, the distance distribution becomes multi-modal, where the number of modes is given by *n*_gen_ + 1. To motivate this effect visually, we fix *N* = 6, *t*_*b*_ = 12, and the per-cell division epimutation rates at 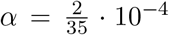 and 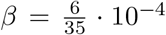. Moreover, we consider *n*_gen_ = 10 branch points, a number that can be assumed for an adult tree. With *t*_end_ = *n*_gen_*t*_*b*_ + *r*, we thus have that a total number of *t*_end_ = 120 + *r* cell divisions occurred along the branching path, where *r* ∈ {1, …, *t*_*b*_}. We consider *r* = 10. Using these fixed values, we evaluate the epimutation process over *L* = 8.6 · 10^6^ CG sites, which corresponds to the number of CG sites in a medium size tree genome.

As can already be seen from (6), the probability distribution of the epignetic distance is a function of *L*. The multi-modality is very conspicuous for large values of *L*. However, in future experimental settings where the distance distribution is determined empirically from single cell DNA methylation data, the number of measured CG sites per cell will be rather sparse (perhaps in the order of only a few percent of the total number of CGs in the genome). To explore the impact of *L* on the probability distribution, we let *L* vary from 1%, 50%, to 100% of the total number of CGs (see **Fig. 4)**. Moreover, we vary *M* from 1 to 6 and study its impact on the epigenetic distance distribution for *L* = 8.6 · 10^6^ (see **Fig. 4A**), *L* = 0.5 · 8.6 · 10^6^ (see **Fig. 4B**), and *L* = 0.01 · 8.6 · 10^6^ (see **Fig. 4C**). The multi-modality that emerges with moderate drift can be seen in all three cases. The position of the modes of the uni-modal distributions are given by ⌊(*L* + 1)*p*_*n*_⌋, which increases for larger values of *n*. This implies that, beginning with the leftmost, the modes correspond to *n* = 1, *n* = 2, …, *n* = *n*_gen_, and *n* = *n*_gen_ + 1, respectively. Moreover, while *M* has an influence on the probability mass function evaluated at the modes, it does not influence the position of the modes. Furthermore, for *L* = 8.6 · 10^6^ and *L* = 0.5 · 8.6 · 10^6^, we observe that the individual uni-modal distributions of the multi-modal distribution for intermediate values of *M* are well separated from each other. However, for *L* = 0.01 · 8.6 · 10^6^, they strongly overlap. Consequently, we find that the different modes become less visible for decreasing values of *L*. From a data analysis standpoint, this could have important implications: If one were to estimate the model parameters, such as *M*, from the empirical distribution using likelihood methods, there would be much less information about these parameters in situations where *L* is small as the multinomial mixture components are less separated from each other (McLachlan and Peel, 2000).

**Figure 4:**
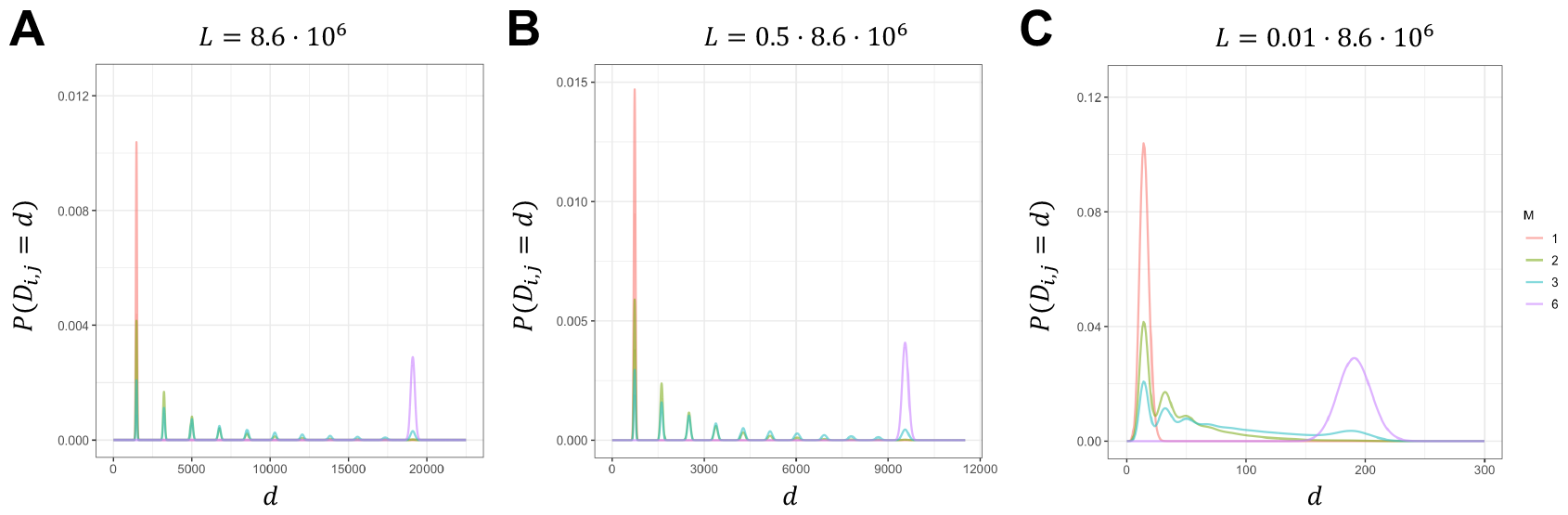
Impact of somatic drift on cell-to-cell epigenetic distance: Distance distribution for different values of *M*, which describes the somatic drift. In all three cases, we assume the following fixed parameter settings: 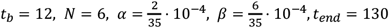 cell divisions. For *M* = 1 (maximal drift, red) and *M* = *N* (no drift, purple), the epigenetic distance consists of a single Binomial distribution while for intermediate *M* values (green and blue), the distance distribution is a multi-modal distribution with *n*_*gen*,_ *+* 1 (here: 11) modes **A**. We consider *L* = 8.6 ⋅ 10^6^ and observe that for intermediate values of *M*, the uni-modal distributions of the multi-modal distribution are well separated. **B**. We consider *L* = 0.5 ⋅ 8.6 ⋅ 10^6^ and observe that also in this case, the uni-modal distributions of the multi-modal distribution are well separated for intermediate values of *M*. **C**. We consider *L* = 0.01 ⋅ 8.6 ⋅ 10^6^ and observe that in this case, especially for larger values of *n*, the uni-modal distributions of the multi-modal distribution overlap for intermediate values of *M*.

Another way to evaluate the impact of drift is by quantifying its effect on the limiting state of the expectation and variance of the epigenetic distance distribution for *n*_gen_ → ∞ (**Fig. 5**). This will be discussed in more detail in the next section. It is intuitive that the limiting state of the expectation of this distribution is higher for *M* = *N* than for *M* = 1, because the cell lineages have had *t*_end_ cell divisions time to diverge from each other, whereas they only have *r* ∈ {1, …, *t*_*b*_} cell division time to diverge when *M* = 1. However, the situation for the limiting state for the variance of the distance distribution is more complex. In (21), we see that for a fixed set of parameter values *α, β, t*_*b*_, *r*, and 1 *< M < N*, we can compute the corresponding 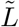 such that for 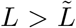, the limit of the variance is smallest for *M* = 1 followed by *M* = *N*, but gets larger for 1 *< M < N*. The reason for this increase is the appearance of additional cell phylogenies that differ from each other in terms of the time to their most recent common ancestor cell (see **Fig. 3**). This source of variation is reflected in the strong multi-modal nature of the distance distribution for intermediate values of *M*. By contrast, for the extremes *M* = 1 and *M* = *N*, the time to the most recent common ancestor cell is uniform across all cell phylogenies, so that the distance distribution consists of only one single Binomial distribution (see **Fig. 4A** and **Fig. 4B**). Interestingly, this ordering does not hold for 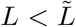. Instead, we find that the limit of the variance increases with increasing *M*. One possible reason, which we can see in **Fig. 5**, is that for small values of *L*, the overlap of the individual Binomial distributions becomes more visible such that the overall variance for intermediate values of *M* is smaller than that for *M* = *N*.

**Figure 5:**
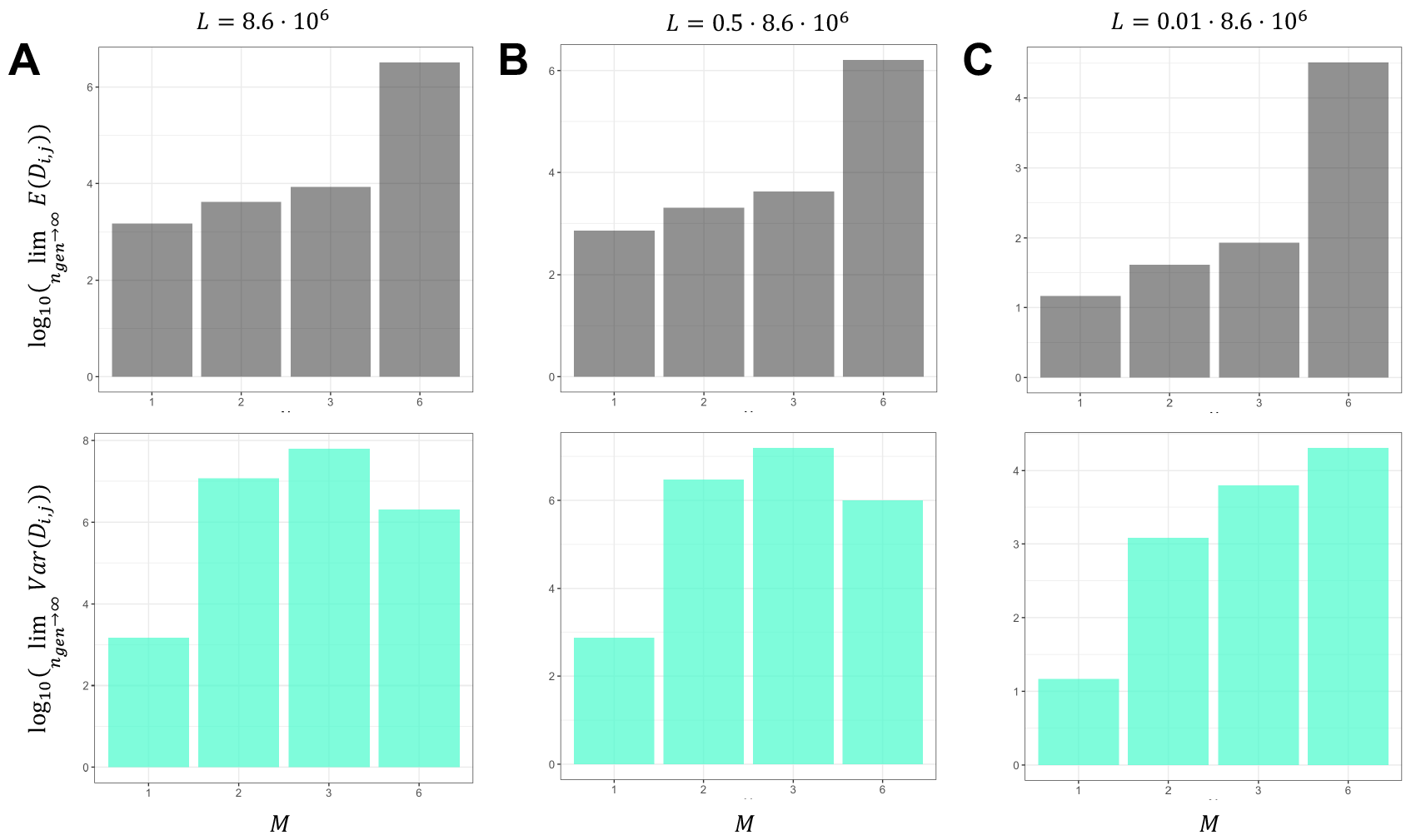
Impact of somatic drift and genome size on the limits of expectation and variance of the cell-to-cell epigenetic distance for increasing number of branch points: In all cases, in gray (turquoise) we have the log_10_-transformation of the limiting states of the expectation (variance) of the distance distribution for *n*_*gen*_ → ∞ for different values of *M* (drift) and *L* (genome size). We assume the following set of parameters: 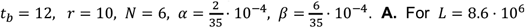. **A**. For *L* = 8.6 ⋅ 10^6^, we observe that the limiting state for the expectation increases with increasing *M* = 3. **B**. For *L* = 0.5 ⋅ 8.6 ⋅ 10^6^, we observe that the limiting state. The limiting state for the variance is the largest for for the expectation increases with increasing *M*. The limiting state for the variance is the largest for *M* = 3. **C**. For *L* = 0.01 ⋅ 8.6 ⋅ 10^6^, we observe that the limiting state for the expectation increases with increasing *M*. However, the limiting state for the variance is the largest for *M* = *N* = 6.

### 3.2 Cell-to-cell DNA methylation heterogeneity converges during successive branching

As we have seen earlier, we can express the developmental time (*t*_end_) in terms of the number of branch points (*n*_gen_). In the following, we explore how the expectation and the variance of the distance distribution converge as *n*_gen_ → ∞. With *t*_end_ = *n*_gen_*t*_*b*_ + *r*, we can write (5) as

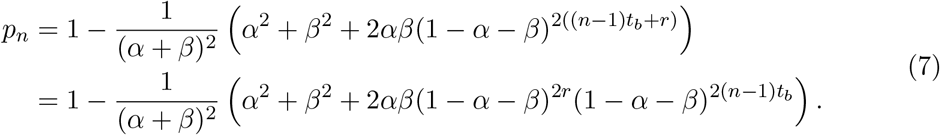

With maximal drift (i.e. *M* = 1) the expectation and the variance of *D*_*i,j*_ are independent of the parameter *n*_gen_. They are given by

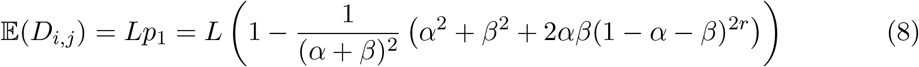

and

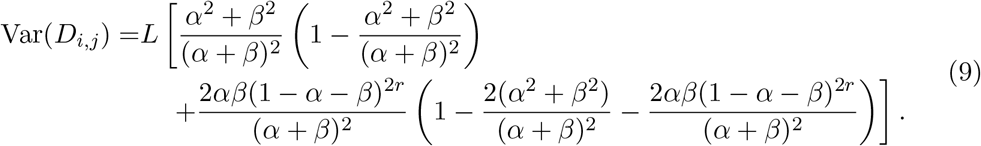

For the extreme case of no drift (i.e. *M* = *N*), we find the expectation of *D*_*i,j*_ for *n*_gen_ → ∞ to be given by

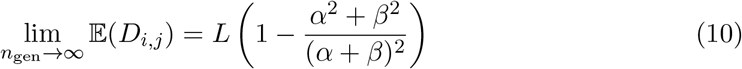

while the variance for *n*_gen_ → ∞ is given by

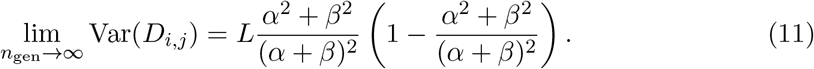

Finally, for moderate drift (i.e. 1 *< M < N*), the expectation of *D*_*i,j*_ for *n*_gen_ → ∞ is expressed by

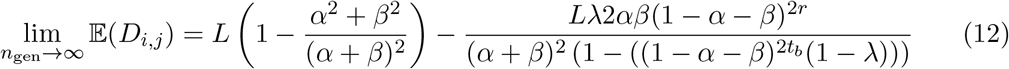

and the variance for *n*_gen_ → ∞ is given by

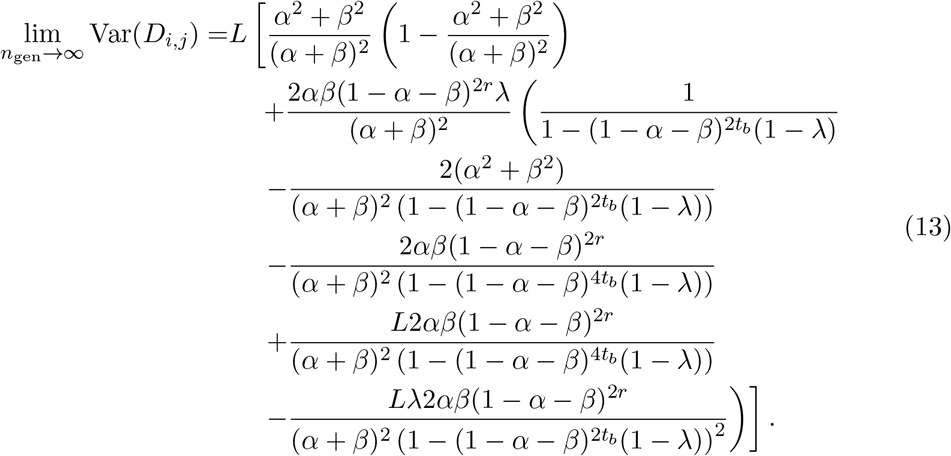

These equations show that as *n*_gen_ → ∞, the expectation and variance of the distance distribution converge to some non-trivial states that are essentially defined by the drift parameter *M* as well as by the epimutation rates *α* and *β* (a detailed derivation is given in Appendix). We refer to these two limits as 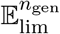 and 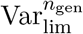 for the expectation and the variance, respectively. We keep in mind that these two values depend on *α, β, r, t*_*b*_, *N, M*, and *L*.

An important empirical question is whether the expectation/variance of the cell-to-cell DNA methylation heterogeneity within the ASC population is far away from, or close to its limiting state value 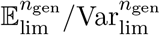 for realistic developmental scenarios. In modular clonal species that use branching for reproduction, the number of branch points can be rather large. Recent age dating of the clonal seagrass *Zostera marina*, for instance, suggests that some clones can be as old as 2000 years (Yu et al., 2023). With about eight branch points per year, the parameter *n*_gen_ can be in the order of tens of thousands. In trees, on the other hand, the life-time number of branching points of an individual is only around 10-20 (Schoen and Schultz, 2019). Setting *n*_gen_ = 10, *L* = 8.6 · 10^6^ and fixing the epimutation rates at 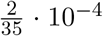 and 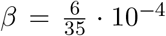, we find that the expected cell-to-cell epigenetic distance is only approximately 0.55% of its limiting state value when *M* = *N*, while for 1 *< M < N* the expected value is approximately between 89.54% and 99.43% of its limiting state. This phenomenon is illustrated in **Fig. 6**, where we keep in mind that the expectation and variance in **Fig. 6** are log_10_-transformed. In addition, we observe that for *M* = *N*, the variance is only approximately 0.88% of the limiting state for *n*_gen_ → ∞, while for 1 *< M < N*, it is approximately between 54.20% and 94.95%. This observation has at least two implications: First, DNA methylation heterogeneity in the ASC population and its derived tissues increases with tree age and may or may not reach saturation depending on the value of *M* and the epimutation rates; and second, differences in developmental rate between individuals (that is, differences in the number of branching points per unit of absolute time) should translate into differences in DNA methylation heterogeneity.

**Figure 6:**
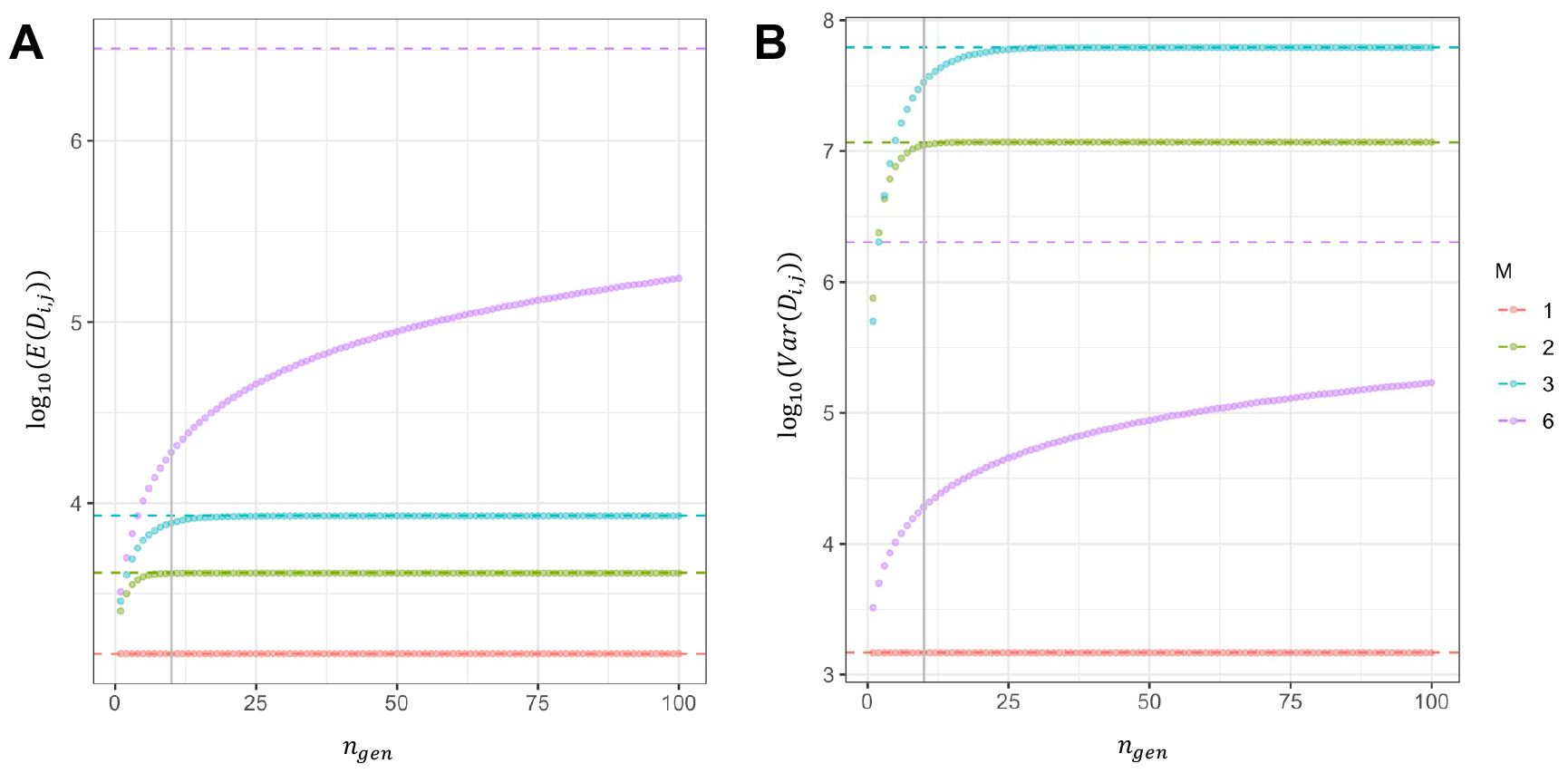
Impact of somatic drift on the limits of expectation and variance of the cell-to-cell epigenetic distance for increasing number of branch points: We assume the following set of parameters: 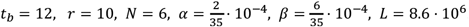. Dashed lines are the limiting states as the number of branch points increases. **A**. We have the log_10_-transformation of the limiting states of the expectation of the distance distribution for *n*_*gen*_ → ∞ for different values of *M*. At *n*_*gen*_ = 10 (gray line), we observe that the expectation for *M* = 2 and *M* = 3 (intermediate drift) is close to its limiting value, while for *M* = *N* (no drift), the expected value is still far away from its limiting state as the number of branch points increases. **B**. We have the log_10_-transformation of the limiting states of the variance of the distance distribution for *n*_*gen*_ → ∞ for different values of *M*. At *n*_*gen*_ = 10 (gray line), we observe that the variance for *M* = 2 and *M* = 3 (no drift) is close to its limiting value, while for *M* = *N* (no drift), the variance is still far away from its limiting state for increasing number of branch points *n*_*gen*_.

**Figure 7:**
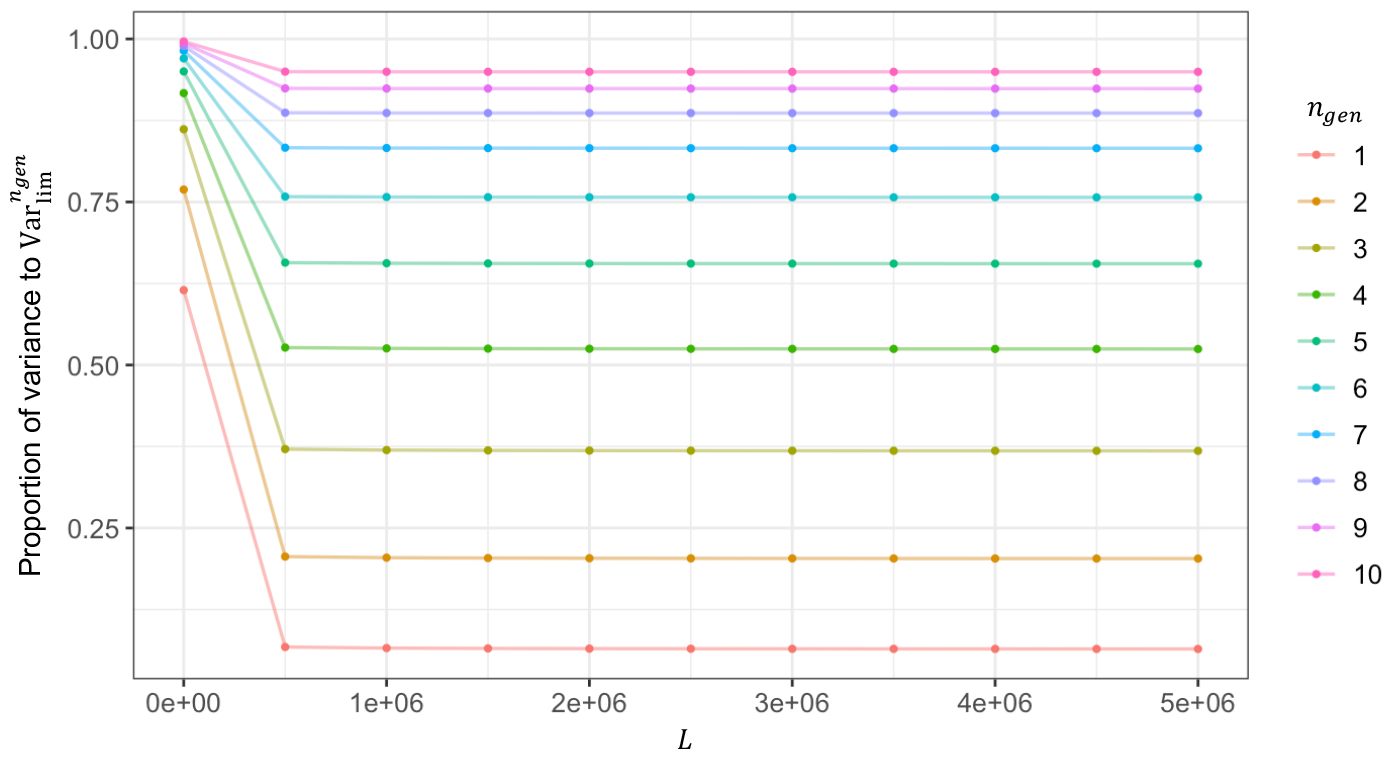
Convergence of proportion of the variance of the cell-to-cell epigenetic distance to 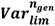 for large values of *L*: We assume the following set of parameters: 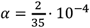. For 11 different values of *L* (points), we computed the corresponding proportion of the variance of *D*_*i,j*_ to 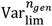. For *n*_*gen*_ = 10, the limit of the proportion for *L* → ∞ is approximately 0.9495.

Another phenomenon is that for a fixed value of *n*_gen_, and letting *α, β, t*_*b*_, *N, M*, and *r* fixed, the proportion of the expectation of the cell-to-cell distance to 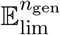 does not depend on *L* (number of CG sites). We can observe the same behavior for the variance when *M* = 1 and *M* = *N*. However, for intermediate values of *M*, the proportion of the variance of the cell-to-cell epigenetic distance to 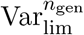 does depend on *L*. Nevertheless, as we can see in (22) in the Appendix, this converges to some value between 0 and 1 for *L* → ∞.

## 4 Discussion

The initiation of new lateral shoots is an important aspect of development and clonal reproduction in plants. Empirical evidence currently supports a “detached meristem model” as the cellular foundation for lateral shoot formation (Wang and Jiao, 2018). According to this model, a few cells are selected from the periphery of the shoot apical meristem (SAM) to serve as precursors for axillary buds and subsequent shoots. The iterative process of branching results in a series of cellular bottlenecks (i.e. somatic drift) that affects the transmission of *de novo* genetic and epigenetic mutations throughout the plant shoot architecture during development. Somatic drift is expected to be particularly relevant for stochastic DNA methylation gains and losses (i.e. spontaneous epimutations) as they are much more frequent than genetic mutations.

In this study, we have examined a special case of the “detached meristem model”, in which precursor cells are randomly selected from the SAM periphery in a way that ensures the independence of cell lineages between two sequential branch points. Under this model, repeated branching leads to a complex mixture of cellular phylogenies that shape the evolution of cell-to-cell DNA methylation heterogeneity within the SAM over time. This process is dependent on the number of branch points, the strength of somatic drift, and the epimutation rate.

The presence of cell-to-cell DNA methylation heterogeneity in the SAM will be a source of epigenetic variation in tissues derived from it. This has direct implications for determining the methylation status of individual cytosines from bulk sequencing data obtained from such tissues, as the read “signal” will derive from a heterogeneous mix of cells. Our work shows that this problem cannot be fully alleviated through layer-specific bulk sampling as the heterogeneity is layer-specific. It remains to be seen if this type of measurement uncertainty has important consequences for recently developed DNA methylation-based phylogenetic inferences approaches (Yao et al., 2023; Yao et al., 2021), particularly in clonal species.

That said, our model could be compatible with emerging single-cell DNA methylation data in plants, such as from single-cell whole genome bisulfite sequencing (scWGBS). Although there are a number of difficult technical challenges in obtaining high quality scWGBS from small cell populations, this approach could provide a statistical framework to pin-point specific model assumptions that are most consistent with observed cell-to-cell DNA methylation patterns. Application of our model to scWGBS could, in principle, yield statistical estimate of the strength of somatic drift (via *M*) and aid in cell lineage reconstruction, both of which are difficult to address with alternative methods.

Our model made several strong assumptions about the independence of cell lineages between two sequential branch points. For instance, we assumed that the mitotic divisions of the apical stem cells are asymmetrical and free of turnover. Also, the sampling of precursor cells along the SAM periphery for *M >* 1 was not spatially confined within clonal sectors of the SAM (see **Fig. 2**). It is known that the axillary meristems are formed at the periphery and tend to draw on several neighboring SAM cells, either within the same clonal sector or straddling sector boundaries. The spatial details about this process depend on the particular plant species and shoot patterning processes, such as phyllotaxy (Kuhlemeier, 2017). Tomimoto and Satake, 2023 recently proposed a more general cell biological modelling approach that accounts for local spatial constraints during precursor sampling. Future work should integrate these spatial constraints with the epimutational models developed here. Nonetheless, the insights gained from our extreme model are useful for gauging the upper bound of the amount of epigenetic heterogeneity that can arise within the SAM. The inclusion of spatial constraints as well as processes such as cell turnover and cell layer invasion will only decrease the expected DNA methylation heterogeneity in the SAM.

## A Derivations

### A.1 Probability distribution of time to MRCA of two apical stem cells

For the derivation of (1)-(3), let *i*_*f*1_ and *j*_*f*1_ be the parent cells of *i* and *j*, respectively and let *i*_*fκ*_ and *j*_*fκ*_ be the parent cells of *i*_*fκ−*1_ and *j*_*fκ−*1_, respectively, where *κ* ∈ {2, …, *n*_gen_ + 1}. We set 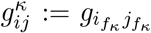. We first note that *g*_*ij*_ = 1 conditioning on *I ≠j* implies that 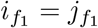, which means that cells *i* and *j* are two different of the *k* identical cells derived from the same precursor cell at the last branch point. Since there are *M* such precursor cells, there are ^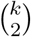^ · *M* possible ways to select *i* and *j*. Therefore, we have

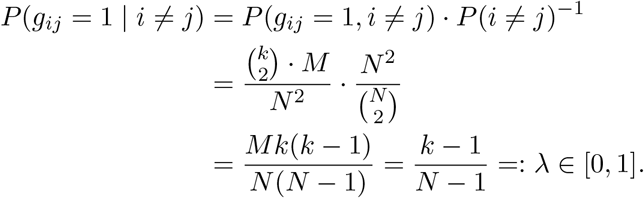

For (2) it holds

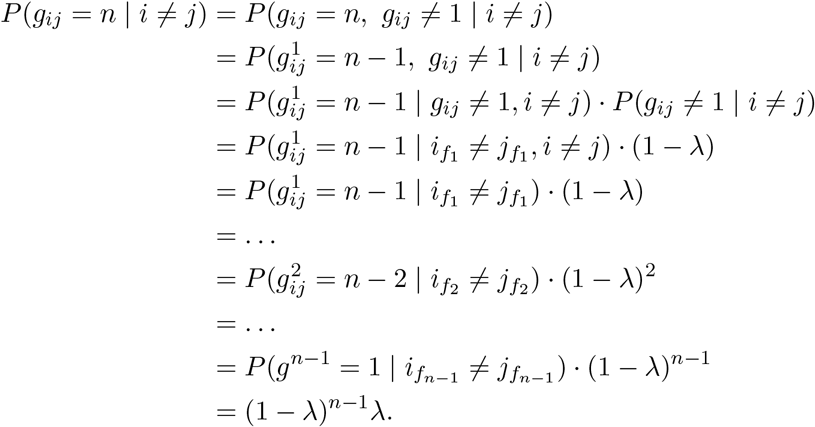

For *n* = *n*_gen_ + 1, note that all cell populations at time *t* ∈ {1, …, *t*_*b*_} originate from the same cell population at *t* = 0 and thus, 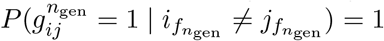, which results in

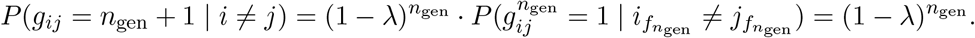

### A.2 Evolution of cell-to-cell DNA methylation divergence

For the derivation of (5), we first note that

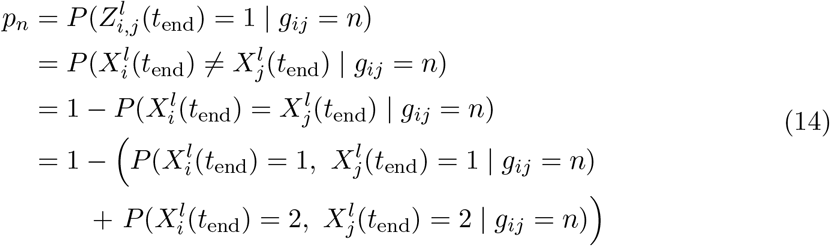

Further, we have

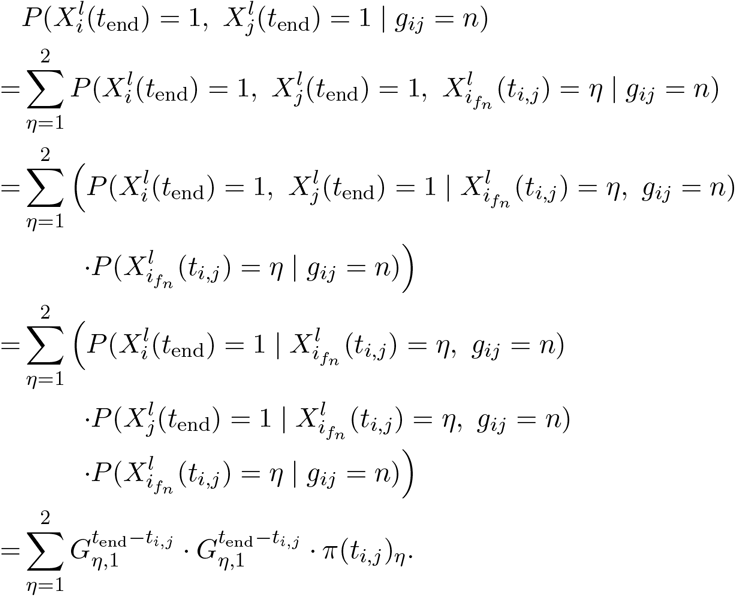

Analogously, we have

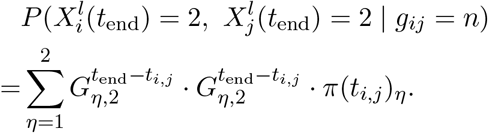

Thus, for Equation 14, we have

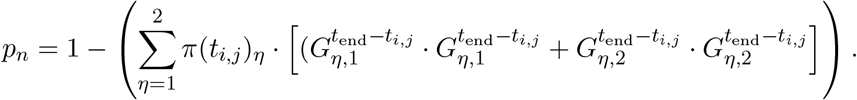

For our model, we further assume that the initial distribution of the methylation states of the loci is the stationary distribution and thus, with *t*_*i,j*_ = (*n*_gen_ + 1 − *n*)*t*_*b*_, we have

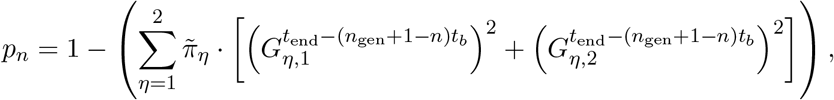

where 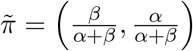 is the stationary distribution of the haploid model as specified in Section 2.3.2. Using the expression given in (4), we have

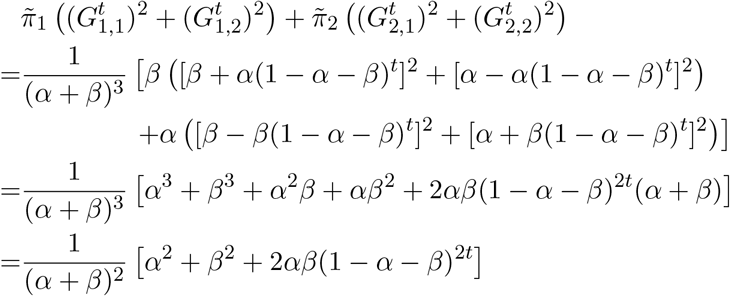

and thus,

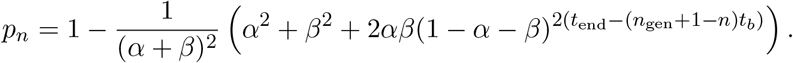

#### A.2.1 Expectation and variance

Using the law of total expectation, the mean of *D*_*i,j*_ is given by

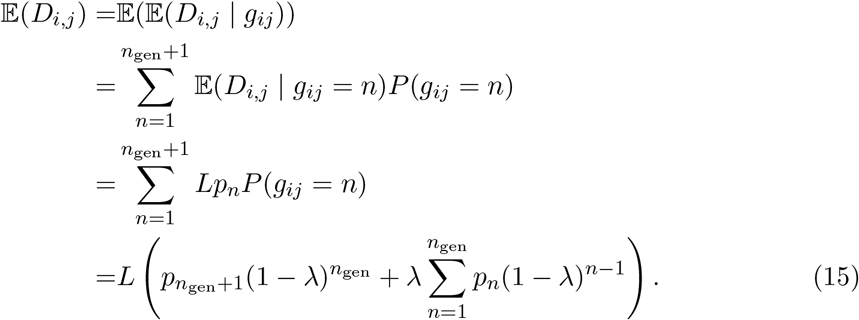

In case of *M* = 1, we have that *λ* = 1 and thus, (15) reduces to (8), which is given by

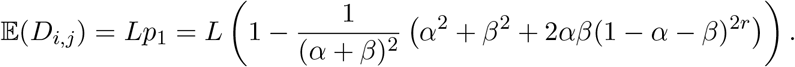

For *M* = *N*, which implies *λ* = 0, we have

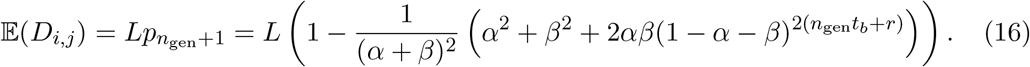

For *n*_gen_ → ∞, we thus have (10), which is given by

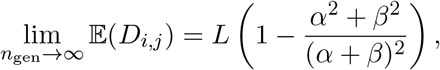

since we assumed *α* and *β* to be sufficiently small.

For *λ* ∈ (0, 1), we observe that

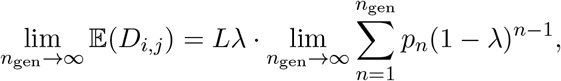

where

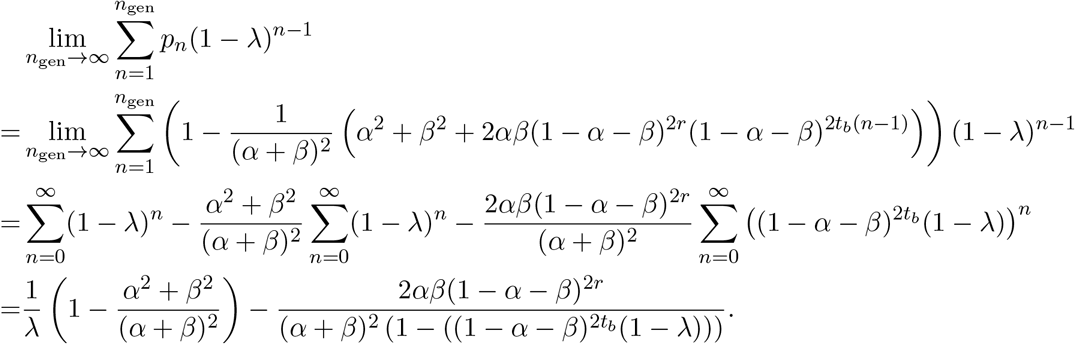

Thus, we have the expression in (12), given by

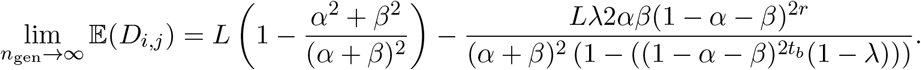

We observe that for *n*_gen_ → ∞, the maximum is obtained for *M* = *N* and since 1−*α*−*β* ∈ (0, 1) we have that 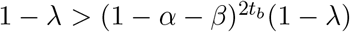 and thus, for *n*_gen_ → ∞, the minimum is obained for *M* = 1.

Now coming to the variance, by applying the law of total variance, we have for the variance of *D*_*i,j*_

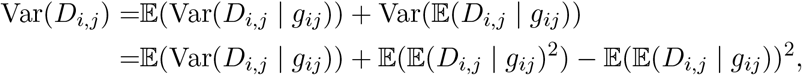

where

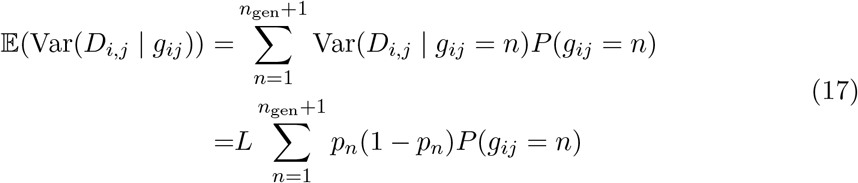

and

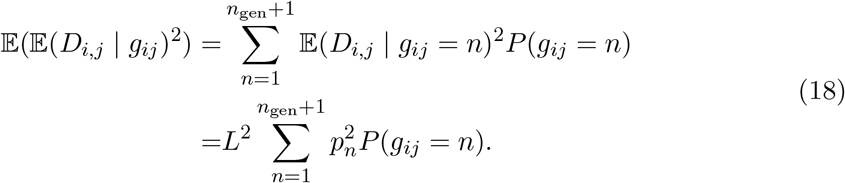

For *M* = 1 and *M* = *N*, we have 𝔼(𝔼(*D*_*i,j*_ | *g*_*ij*_)^2^) = 𝔼(𝔼(*D*_*i,j*_ | *g*_*ij*_))^2^ and thus,

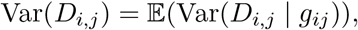

which is given by *Lp*_1_(1 − *p*_1_) and *Lp*_*n*_gen+1(1 − *p*_*n*_gen+1), respectively. With

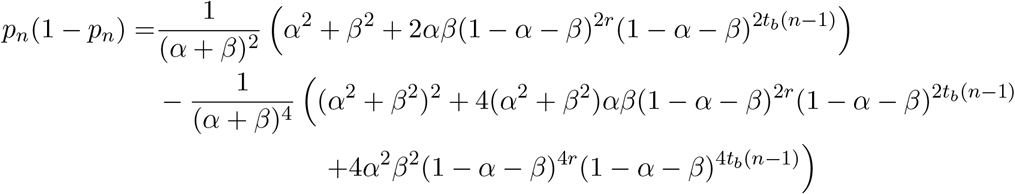

and

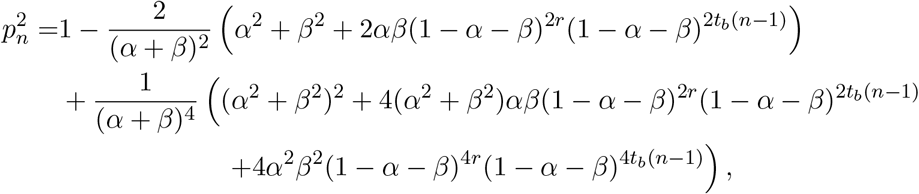

we have in case of *M* = 1 the expression in (9), which is given by

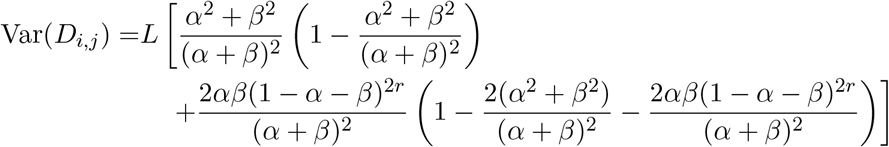

Note that with

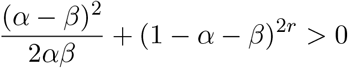

we have

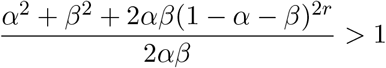

and thus it holds that

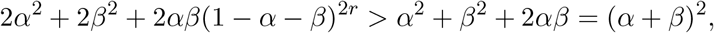

which yields that

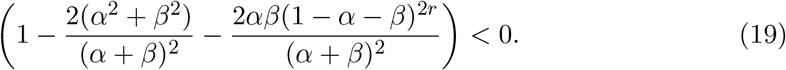

In case of *M* = *N* the expression in (11), which is given by

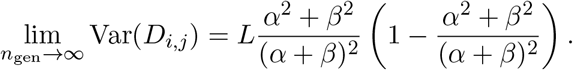

From (19), we observe that the variance for *M* = 1 is always smaller than the the variance for *M* = *N* for *n*_gen_ → ∞.

For 1 *< M < N*, we have *λ* ∈ (0, 1) and in a similar way as we have computed 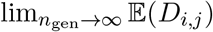, we can show that

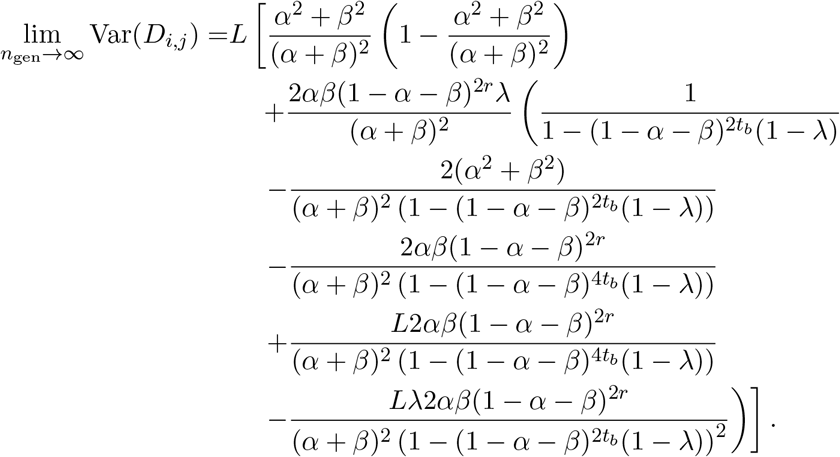

We observe that for *L* = 1, the sum in the last bracket becomes

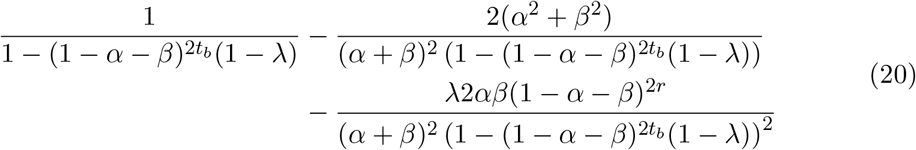

With

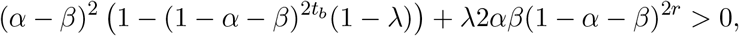

which is equivalent to

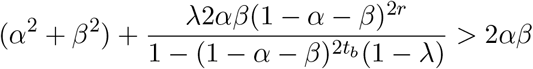

and thus, we have

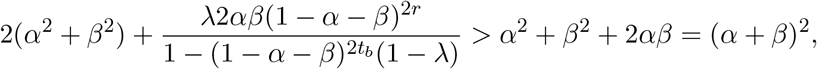

which yields

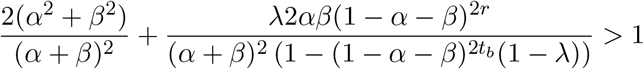

It follows that (20) is negative, which means that for small values of *L* and *n*_gen_ → ∞, the variance of *D*_*i,j*_ for 1 *< M < N* is smaller than that for *M* = *N*.

We note that

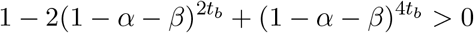

is always satisfied and with *λ* ∈ (0, 1), we have

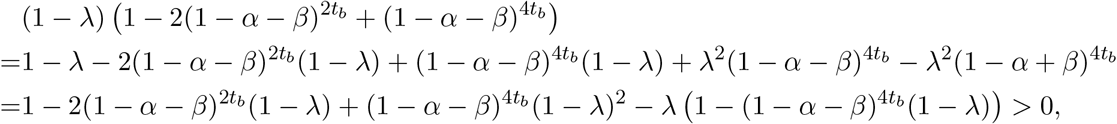

which yields

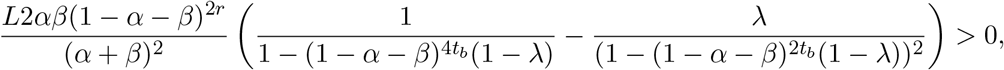

since *α, β >* 0. Thus, as *L* increases, for *n*_gen_ → ∞, the variance of *D*_*i,j*_ for 1 *< M < N* eventually becomes larger than that for *M* = *N*. In particular, for a fixed 1 *< M < N*, this occurs at

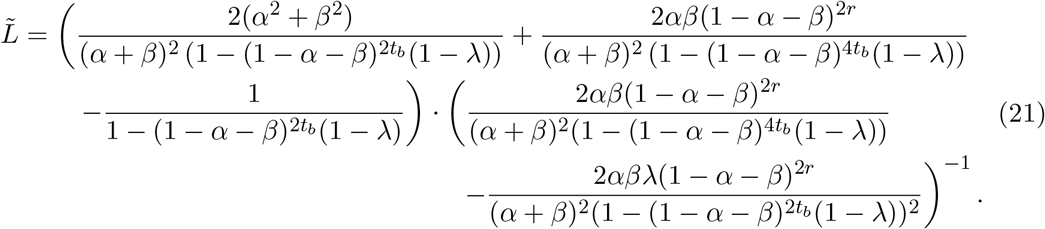

Let 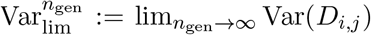 and 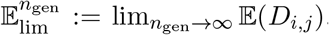. The proportion of the variance of the cell-to-cell distance and its limit for *n*_gen_ → ∞ for some 1 *< M < N*, i.e. 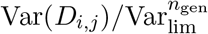 is given by

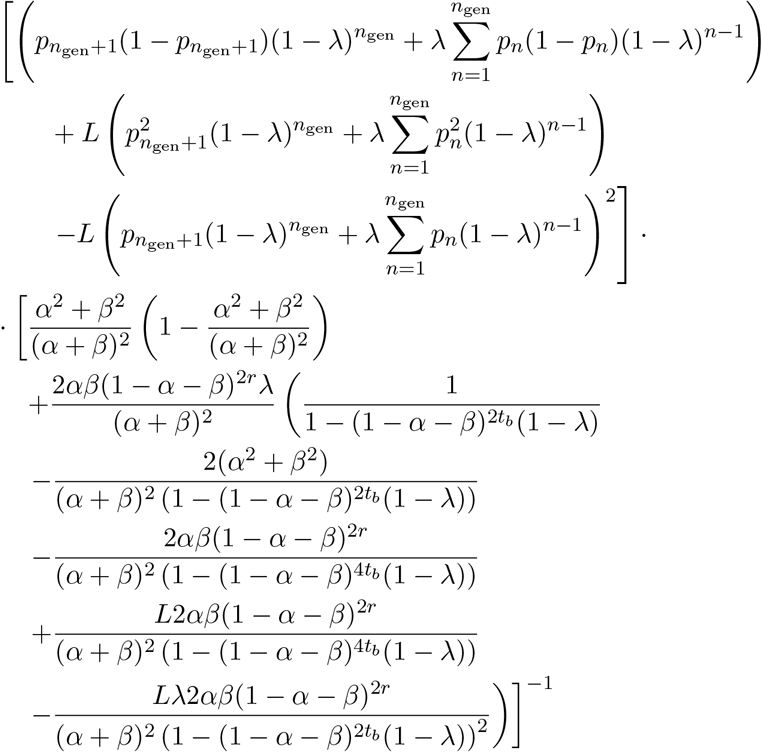

For *L* → ∞, L’Hospital’s Rule yields

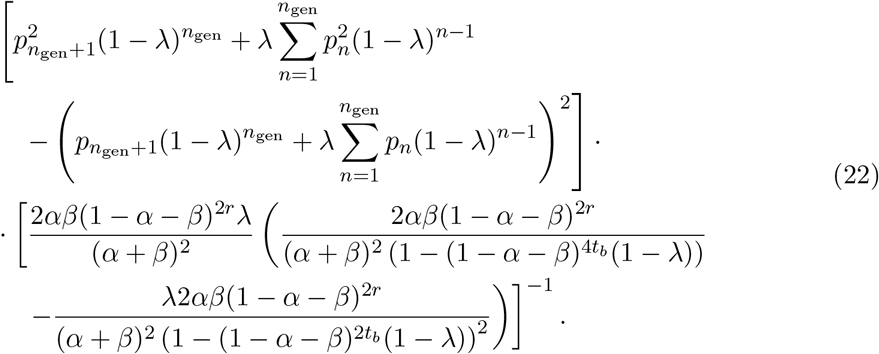

## References

Antolin, M. F., & Strobeck, C. (1985). The population genetics of somatic mutation in plants. The American Naturalist, 126 (1), 52–62. 10.1086/284395

Barbier de Reuille, P., Routier-Kierzkowska, A.-L., Kierzkowski, D., Bassel, G. W., Schüpbach, T., Tauriello, G., Bajpai, N., Strauss, S., Weber, A., Kiss, A., Burian, A., Hofhuis, H., Sapala, A., Lipowczan, M., Heimlicher, M. B., Robinson, S., Bayer, E. M., Basler, K., Koumoutsakos, P., … Smith, R. S. (2015). MorphoGraphX: A platform for quantifying morphogenesis in 4d (D. C. Bergmann, Ed.). eLife, 4, e05864. 10.7554/eLife.05864

Berger, F., & Twell, D. (2011). Germline specification and function in plants. Annual Review of Plant Biology, 62 (1), 461–484. 10.1146/annurev-arplant-042110-103824

Briffa, A., Hollwey, E., Shahzad, Z., Moore, J. D., Lyons, D. B., Howard, M., & Zilberman, D. (2023). Millennia-long epigenetic fluctuations generate intragenic DNA methylation variance in arabidopsis populations. Cell Systems, 14 (11), 953–967.e17. 10.1016/j.cels.2023.10.007

Burian, A. (2021). Does shoot apical meristem function as the germline in safeguarding against excess of mutations? Frontiers in Plant Science, 12. Retrieved January 5, 2024, from https://www.frontiersin.org/articles/10.3389/fpls.2021.707740

Burian, A., Barbier de Reuille, P., & Kuhlemeier, C. (2016). Patterns of stem cell divisions contribute to plant longevity. Current Biology, 26 (11), 1385–1394. 10.1016/j.cub.2016.03.067

Domagalska, M. A., & Leyser, O. (2011). Signal integration in the control of shoot branching. Nature Reviews Molecular Cell Biology, 12 (4), 211–221. 10.1038/nrm3088

Folse, H. J., III, & Roughgarden, J. (2012). DIRECT BENEFITS OF GENETIC MOSAICISM AND INTRAORGANISMAL SELECTION: MODELING COEVOLUTION BETWEEN a LONG-LIVED TREE AND a SHORT-LIVED HERBIVORE. Evolution, 66 (4), 1091–1113. 10.1111/j.1558-5646.2011.01500.x

Foster, T. M., & Aranzana, M. J. (2018). Attention sports fans! the far-reaching contributions of bud sport mutants to horticulture and plant biology. Horticulture Research, 5 (1), 1–13. 10.1038/s41438-018-0062-x

Frank, M. H., & Chitwood, D. H. (2016). Plant chimeras: The good, the bad, and the ‘bizzaria’. Developmental Biology, 419 (1), 41–53. 10.1016/j.ydbio.2016.07.003

Heidstra, R., & Sabatini, S. (2014). Plant and animal stem cells: Similar yet different. Nature Reviews Molecular Cell Biology, 15 (5), 301–312. 10.1038/nrm3790

Herrera, C. M., Bazaga, P., Pérez, R., & Alonso, C. (2021). Lifetime genealogical divergence within plants leads to epigenetic mosaicism in the shrub lavandula latifolia (lamiaceae). New Phytologist, 231 (5), 2065–2076. 10.1111/nph.17257

Hofmeister, B. T., Denkena, J., Colomé-Tatché, M., Shahryary, Y., Hazarika, R., Grimwood, J., Mamidi, S., Jenkins, J., Grabowski, P. P., Sreedasyam, A., Shu, S., Barry, K., Lail, K., Adam, C., Lipzen, A., Sorek, R., Kudrna, D., Talag, J., Wing, R., … Schmitz, R. J. (2020). A genome assembly and the somatic genetic and epigenetic mutation rate in a wild long-lived perennial populus trichocarpa. Genome Biology, 21 (1), 259. 10.1186/s13059-020-02162-5

Iwasa, Y., Tomimoto, S., & Satake, A. (2023). The genetic structure within a single tree is determined by the behavior of the stem cells in the meristem. Genetics, 223 (4), iyad020. 10.1093/genetics/iyad020

Jackson, M. D. B., Duran-Nebreda, S., Kierzkowski, D., Strauss, S., Xu, H., Landrein, B., Hamant, O., Smith, R. S., Johnston, I. G., & Bassel, G. W. (2019). Global topological order emerges through local mechanical control of cell divisions in the arabidopsis shoot apical meristem. Cell Systems, 8 (1), 53–65.e3. 10.1016/j.cels.2018.12.009

Johannes, F., & Schmitz, R. J. (2019). Spontaneous epimutations in plants. New Phytologist, 221 (3), 1253–1259. 10.1111/nph.15434

Klekowski, E. J., & Kazarinova-Fukshansky, N. (1984a). Shoot apical meristems and mutation: Fixation of selectively neutral cell genotypes. American Journal of Botany, 71 (1), 22–27. 10.2307/2443619

Klekowski, E. J., & Kazarinova-Fukshansky, N. (1984b). Shoot apical meristems and mutation: Selective loss of disadvantageous cell genotypes. American Journal of Botany, 71 (1), 28–34. 10.2307/2443620

Klekowski, E. J., Kazarinova-Fukshansky, N., & Fukshansky, L. (1989). Patterns of plant ontogeny that may influence genomic stasis. American Journal of Botany, 76 (2), 185–195. 10.2307/2444660

Klekowski, E. J., Kazarinova-Fukshansky, N., & Mohr, H. (1985). Shoot apical meristems and mutation: Stratified meristems and angiosperm evolution. American Journal of Botany, 72 (11), 1788–1800. 10.2307/2443736

Kuhlemeier, C. (2017). Phyllotaxis. Current Biology, 27 (17), R882–R887. 10.1016/j.cub.2017.05.069

Kwiatkowska, D. (2008). Flowering and apical meristem growth dynamics. Journal of Experimental Botany, 59 (2), 187–201. 10.1093/jxb/erm290

Laufs, P., Grandjean, O., Jonak, C., Kîeu, K., & Traas, J. (1998). Cellular parameters of the shoot apical meristem in arabidopsis. The Plant Cell, 10 (8), 1375–1389. 10.1105/tpc.10.8.1375

Lyndon, R. F. (1998, May 28). The shoot apical meristem: Its growth and development. Cambridge University Press.

McLachlan, G. J., & Peel, D. (2000, October 2). Finite mixture models. John Wiley & Sons.

McSteen, P., & Leyser, O. (2005). Shoot branching. Annual Review of Plant Biology, 56 (1), 353–374. 10.1146/annurev.arplant.56.032604.144122

Nicolas, A., & Laufs, P. (2022). Meristem initiation and de novo stem cell formation. Frontiers in Plant Science, 13. Retrieved January 16, 2024, from https://www.frontiersin.org/articles/10.3389/fpls.2022.891228

Otto, S. P., & Orive, M. E. (1995). Evolutionary consequences of mutation and selection within an individual. Genetics, 141 (3), 1173–1187. 10.1093/genetics/141.3.1173

Paszkowski, J., & Grossniklaus, U. (2011). Selected aspects of transgenerational epigenetic inheritance and resetting in plants. Current Opinion in Plant Biology, 14 (2), 195–203. 10.1016/j.pbi.2011.01.002

Pineda-Krch & Fagerström. (1999). On the potential for evolutionary change in meristematic cell lineages through intraorganismal selection. Journal of Evolutionary Biology, 12 (4), 681–688. 10.1046/j.1420-9101.1999.00066.x

Pineda-krch, M., & Lehtilä, K. (2002). Cell lineage dynamics in stratified shoot apical meristems. Journal of Theoretical Biology, 219 (4), 495–505. 10.1006/jtbi.2002.3139

Plomion, C., Aury, J.-M., Amselem, J., Leroy, T., Murat, F., Duplessis, S., Faye, S., Francillonne, N., Labadie, K., Le Provost, G., Lesur, I., Bartholomé, J., Faivre-Rampant, P., Kohler, A., Leplé, J.-C., Chantret, N., Chen, J., Diévart, A., Alaeitabar, T., … Salse, J. (2018). Oak genome reveals facets of long lifespan. Nature Plants, 4 (7), 440–452. 10.1038/s41477-018-0172-3

Reddy, G. V., Heisler, M. G., Ehrhardt, D. W., & Meyerowitz, E. M. (2004). Real-time lineage analysis reveals oriented cell divisions associated with morphogenesis at the shoot apex of arabidopsis thaliana. Development, 131 (17), 4225–4237. 10.1242/dev.01261

Reusch, T. B. H., Baums, I. B., & Werner, B. (2021). Evolution via somatic genetic variation in modular species. Trends in Ecology & Evolution, 36 (12), 1083–1092. 10.1016/j.tree.2021.08.011

Romberger, J., Romberger, J., Hejnowicz, Z., & Hill, J. (2004). Plant structure: Function and development : A treatise on anatomy and vegetative development, with special reference to woody plants. Blackburn Press. https://books.google.de/books?id=yB9TAAAACAAJ

Satake, A., Imai, R., Fujino, T., Tomimoto, S., Ohta, K., Na’iem, M., Indrioko, S., Widiyatno Purnomo, S., Mollá-Morales, A., Nizhynska, V., Tani, N., Suyama, Y., Sasaki, E., & Kasahara, M. (2023). Somatic mutation rates scale with time not growth rate in long-lived tropical trees. eLife, 12. 10.7554/eLife.88456

Schmid-Siegert, E., Sarkar, N., Iseli, C., Calderon, S., Gouhier-Darimont, C., Chrast, J., Cattaneo, P., Schütz, F., Farinelli, L., Pagni, M., Schneider, M., Voumard, J., Jaboyedoff, M., Fankhauser, C., Hardtke, C. S., Keller, L., Pannell, J. R., Reymond, A., Robinson-Rechavi, M., … Reymond, P. (2017). Low number of fixed somatic mutations in a long-lived oak tree. Nature Plants, 3 (12), 926–929. 10.1038/s41477-017-0066-9

Schoen, D. J., & Schultz, S. T. (2019). Somatic mutation and evolution in plants. Annual Review of Ecology, Evolution, and Systematics, 50 (1), 49–73. 10.1146/annurev-ecolsys-110218-024955

Shahryary, Y., Symeonidi, A., Hazarika, R. R., Denkena, J., Mubeen, T., Hofmeister, B., van Gurp, T., Colomé-Tatché, M., Verhoeven, K. J., Tuskan, G., Schmitz, R. J., & Johannes, F. (2020). AlphaBeta: Computational inference of epimutation rates and spectra from high-throughput DNA methylation data in plants. Genome Biology, 21 (1), 260. 10.1186/s13059-020-02161-6

Shi, B., Zhang, C., Tian, C., Wang, J., Wang, Q., Xu, T., Xu, Y., Ohno, C., Sablowski, R., Heisler, M. G., Theres, K., Wang, Y., & Jiao, Y. (2016). Two-step regulation of a meristematic cell population acting in shoot branching in arabidopsis. PLOS Genetics, 12 (7), e1006168. 10.1371/journal.pgen.1006168

Stewart, R. N., & Dermen, H. (1970). Determination of number and mitotic activity of shoot apical initial cells by analysis of mericlinal chimeras. American Journal of Botany, 57 (7), 816–826. 10.1002/j.1537-2197.1970.tb09877.x

Tomimoto, S., Iwasa, Y., & Satake, A. (2023, October 3). Branching architecture affects genetic diversity within an individual tree. 10.1101/2023.10.02.560431

Tomimoto, S., & Satake, A. (2023). Modelling somatic mutation accumulation and expansion in a long-lived tree with hierarchical modular architecture. Journal of Theoretical Biology, 565, 111465. 10.1016/j.jtbi.2023.111465

van der Graaf, A., Wardenaar, R., Neumann, D. A., Taudt, A., Shaw, R. G., Jansen, R. C., Schmitz, R. J., Colomé-Tatché, M., & Johannes, F. (2015). Rate, spectrum, and evolutionary dynamics of spontaneous epimutations. Proceedings of the National Academy of Sciences, 112 (21), 6676–6681. 10.1073/pnas.1424254112

Wang, Y., & Jiao, Y. (2018). Auxin and above-ground meristems. Journal of Experimental Botany, 69 (2), 147–154. 10.1093/jxb/erx299

Watson, J. M., Platzer, A., Kazda, A., Akimcheva, S., Valuchova, S., Nizhynska, V., Nordborg, M., & Riha, K. (2016). Germline replications and somatic mutation accumulation are independent of vegetative life span in arabidopsis. Proceedings of the National Academy of Sciences, 113 (43), 12226–12231. 10.1073/pnas.1609686113

Whitham, T. G., & Slobodchikoff, C. N. (1981). Evolution by individuals, plant-herbivore interactions, and mosaics of genetic variability: The adaptive significance of somatic mutations in plants. Oecologia, 49 (3), 287–292. 10.1007/BF00347587

Yao, N., Zhang, Z., Yu, L., Hazarika, R., Yu, C., Jang, H., Smith, L. M., Ton, J., Liu, L., Stachowicz, J. J., Reusch, T. B. H., Schmitz, R. J., & Johannes, F. (2023). An evolutionary epigenetic clock in plants. Science, 381 (6665), 1440–1445. 10.1126/science.adh9443

Yao, N., Schmitz, R. J., & Johannes, F. (2021). Epimutations define a fast-ticking molecular clock in plants. Trends in Genetics, 37 (8), 699–710. 10.1016/j.tig.2021.04.010

Yu, L., Boström, C., Franzenburg, S., Bayer, T., Dagan, T., & Reusch, T. B. H. (2020). Somatic genetic drift and multilevel selection in a clonal seagrass. Nature Ecology & Evolution, 4 (7), 952–962. 10.1038/s41559-020-1196-4

Yu, L., Renton, J., Burian, A., Khachaturyan, M., Kotta, J., Stachowicz, J. J., DuBois, K., Baums, I. B., Werner, B., & Reusch, T. B. H. (2023, November 9). Precise age estimation in clonal species using a somatic genetic clock. 10.1101/2023.11.07.566010

